# Stable iPSC-derived NKX2-1^+^ Lung Bud Tip Progenitor Organoids Give Rise to Airway and Alveolar Cell Types

**DOI:** 10.1101/2022.02.25.481981

**Authors:** Renee F.C. Hein, Ansley S. Conchola, Alexis Fine, Zhiwei Xiao, Tristan Frum, Charlie J. Childs, Yu-Hwai Tsai, Emily M. Holloway, Sha Huang, John Mahoney, Jason R. Spence

## Abstract

Bud tip progenitors (BTPs) in the developing lung give rise to all epithelial cell types found in the airways and alveoli. The current work aimed to develop an iPSC organoid model enriched with stable NKX2-1^+^ BTP-like cells. Building on prior work, we optimized a directed differentiation paradigm to generate spheroids with robust NKX2-1 expression. Spheroids were expanded into organoids that possessed NKX2-1^+^/CPM^+^ BTP-like cells, which increased in number over time. Single cell RNA-sequencing analysis revealed a high degree of transcriptional similarity between induced BTPs (iBTPs) and *in vivo* BTPs. Using FACS, iBTPs can be purified and expanded as induced bud tip organoids (iBTO), which maintain an enriched population of bud tip progenitors. When iBTOs are directed to differentiate into airway or alveolar cell types using well-established methods, they give rise to organoids composed of organized airway or alveolar epithelium, respectively. Collectively, iBTOs are transcriptionally and functionally similar to *in vivo* BTPs, providing an important model to study human lung development and differentiation.

**SUMMARY STATEMENT:** iPSC-derived lung bud tip progenitors emerge in organoid culture, can be isolated and expanded, are transcriptionally similar to primary bud tip progenitors, and can differentiate into airway or alveolar organoids.

## INTRODUCTION

Advances in directed differentiation methods have led to the development of numerous embryonic or induced pluripotent stem cell (iPSC)-derived cell or organoid models of the airway and alveoli, which have enhanced our ability to model human lung development and disease (Wang *et al*., 2007; Mou *et al*., 2012; Gotoh *et al*., 2014; Huang *et al*., 2014; Dye *et al*., 2015, 2016; Konishi *et al*., 2016; Chen *et al*., 2017; McCauley *et al*., 2017; Yamamoto *et al*., 2017, 2020; Hawkins *et al*., 2017; Miller *et al*., 2018, 2019; Tamò *et al*., 2018; de Carvalho *et al*., 2019; Jacob *et al*., 2019; Leibel *et al*., 2020). Alveolar and airway cell types in mice and humans are derived from a common, developmentally transient progenitor population, called bud tip progenitors, which reside at the tips of the branching tree-like network of epithelial tubes that make up the lung epithelium (Serra, Pelton and Moses, 1994; Perl *et al*., 2005; Abler, Mansour and Sun, 2008; Goss *et al*., 2009; Rawlins *et al*., 2009; Rockich *et al*., 2013; Alanis *et al*., 2014; Nikolić *et al*., 2017; Miller *et al*., 2018).

Bud tip progenitors obtained from the human fetal lung and grown as organoids serve as a useful tool to study the mechanisms responsible for bud tip progenitor cell maintenance and differentiation into airway and alveolar cell types (Nikolić *et al*., 2017; Miller *et al*., 2018, 2020; Conway *et al*., 2020; Sun, Evans and Rawlins, 2020). Despite this progress, organoids derived from fetal tissue are not broadly accessible to the research community and are associated with ethical and regulatory challenges, emphasizing the importance of iPSC-derived lung models. While we and others have made some progress in developing bud tip progenitor cell-like models from iPSCs (Chen *et al*., 2017; Miller *et al*., 2018), new technologies such as single cell RNA sequencing (scRNA-seq) and single cell lineage tracing have highlighted off-target cell types and unexpected plasticity in iPSC-derived cultures, alongside unexpected plasticity, where cells appear committed to a specific cell type or cell lineage but change fate (Little *et al*., 2019; Hurley *et al*., 2020). This concept of cellular plasticity within the lung has also been demonstrated *in vivo,* where hyperactive WNT signaling in lung progenitors was shown to cause an emergence of intestinal cells in transgenic mouse embryos (Okubo and Hogan, 2004). Therefore, a current challenge in the field, addressed in the current work, is to develop a long-lived and transcriptionally stable bud tip progenitor-like model from iPSCs.

Single cell RNA sequencing technologies have also made it possible to benchmark iPSC-derived cultures against primary tissue to compare transcriptional similarity and to accurately catalogue the diversity of on-target or off-target cell types observed *in vitro* (Hawkins *et al*., 2017, 2021; McCauley *et al*., 2018; Holloway *et al*., 2020; Hor *et al*., 2020; Yu *et al*., 2020). Since iPSC-derived cultures are known to be plastic, and the fact that iPSC differentiation is not 100% efficient, benchmarking has become an important step towards understanding the full complement of cells present in a culture. The current manuscript therefore also sought to benchmark iPSC-derived bud tip progenitor organoids (iBTOs) to interrogate the diversity of cell types in culture and similarity to bud tip progenitor organoids from the fetal lung.

Here, we optimized an iPSC directed differentiation paradigm to generate self-organizing 3D spheroids with robust NKX2-1 expression. Expansion of NKX2-1^+^ cells in bud tip progenitor media over 3 – 17 weeks gives rise to heterogenous organoids that contain NKX2-1^+^ bud tip progenitor-like cells co-expressing markers of human bud tip progenitors, including SOX9, SOX2 and the cell surface marker CPM (Yamamoto *et al*., 2020). Using an NKX2-1 reporter iPSC line along with CPM to quantitatively assess cultures via flow cytometry, we observed that bud tip progenitor-like cells expanded over subsequent weeks in culture. FACS isolation and further culture allowed for the expansion of NKX2-1^+^/CPM^+^ cells as bud tip progenitor-like organoids (iBTOs) that maintained ∼85% NKX2-1^+^/CPM^+^ cells for at least 8 weeks. scRNA-seq analysis of bud tip progenitor cells from unsorted organoids or following FACS-enrichment revealed a high degree of transcriptional similarity to fetal-derived bud tip organoids as well as a shared transcriptional signature with *in vivo* bud tip progenitors. In addition, scRNA-seq from iBTOs that have spent less (3 weeks) or more (10 weeks) time in culture suggest that induced bud tip progenitors become more transcriptionally similar to native bud tip progenitors as they age. Finally, we used well-established methods to direct differentiation of iBTOs into organoids composed of airway epithelium (including basal, secretory, ciliated, goblet, and neuroendocrine cells) or alveolar type II (AT2) organoids (Jacob *et al*., 2017; Miller *et al*., 2020). Collectively, this study describes a robust method to generate stable bud tip progenitor-like cells from iPSCs that closely resemble organoids derived from primary tissue. This model can be readily used to study lung development and illustrates a proof-of-concept for cellular engineering and cell therapy.

## RESULTS

### Lung Spheroids are Optimized for NXK2.1 Expression but Remain Heterogenous

NKX2-1 is the earliest marker during lung epithelial specification (Lazzaro *et al*., 1991) and loss of NKX2-1 leads to lung agenesis (Minoo *et al*., 1995, 1999; Little *et al*., 2019; Kuwahara *et al*., 2020). We therefore sought to build on a previously-published method to generate iPSC-derived foregut spheroids (Dye *et al*., 2015) by optimizing endoderm induction efficiency, foregut spheroid development, and finally to improve conditions that induced robust NKX2-1 expression.

We began by testing conditions to improve definitive endoderm (DE) differentiation efficiency and reproducibility. DE induction of varying efficiencies is achieved by using Activin A (ACTA) ligand (Kubo *et al*., 2004; D’Amour *et al*., 2006; Spence *et al*., 2011) with WNT and BMP signaling playing a synergistic role during the initial stages DE specification (Gadue *et al*., 2006; Green *et al*., 2011; Loh *et al*., 2014; Matsuno *et al*., 2016; Rankin *et al*., 2018; Heemskerk *et al*., 2019). Therefore, we tested combinations of ACTA alongside the small molecule WNT activator CHIR99021 (CHIR) or BMP4 on the first day of a three-day ACTA differentiation culture (Fig. S1A,B). Using flow cytometry with a SOX17-tdTomato and SOX2-mCITRINE hESC reporter line (Martyn *et al*., 2018) to quantitate cell composition, we observed that ACTA alone induced 48.41% SOX17^+^ cells, while the addition of CHIR or BMP both enhanced DE differentiation, leading to 96.48% SOX17^+^ cells or 86.83% SOX17^+^ cells respectively. Addition of both CHIR and BMP4 led to 71.03% SOX17^+^ induction (Fig. S1A). DE cultures from an additional cell line were co-stained with SOX17 and FOXA2 to confirm definitive endoderm cell identity (Fig. S1B). We observed that near-pure SOX17^+^ cultures obtained via ACTA/CHIR failed to give rise to self-organizing 3D spheroids (data not shown), consistent with published data showing that self-organizing foregut and hindgut organoids consist of both epithelium and mesenchymal lineages (Spence *et al*., 2011; Dye *et al*., 2015, 2016). Therefore, ACTA/BMP4 was used for subsequent experiments.

Following DE specification, monolayers were directed into 3D foregut endoderm spheroids by combining a method that efficiently induces ventral foregut endoderm competent to be specified as lung (Rankin *et al*., 2016) alongside methods that induce 3D self-organization (Spence *et al*., 2011; Dye *et al*., 2015). This included BMP inhibition via NOGGIN (NOG) for 3 days plus all trans retinoic acid (ATRA) on the last day as well as FGF4 and CHIR, which are required for 3D spheroid formation, for all 3 days (Fig. S1C,D). After 3 days, spheroids were collected and suspended in Matrigel and treated for 3 additional days with low-BMP4 as well as WNT3A and RSPO1, which together stimulate WNT signaling (Fig. S1E) (Shu *et al*., 2005; Goss *et al*., 2009; Harris-Johnson *et al*., 2009; Domyan *et al*., 2011; Jacobs, Ku and Que, 2012; Miller *et al*., 2012; Serra *et al*., 2017). WNT activation through RSPO/WNT or CHIR resulted in comparable expression levels of foregut and hindgut markers (Fig. S1E); however, CHIR is a GSK3β inhibitor and can have non-WNT mediated effects, so RSPO/WNT were used as more specific activators of WNT signaling.

At the end of the 9-day directed differentiation (Fig. 1A), spheroids were analyzed for *NKX2-1* expression via qRT-PCR (on day 10) (Fig. 1B). When directly compared to spheroids generated using previously published foregut/lung spheroid protocols (Dye *et al*., 2015; Miller *et al*., 2019), NKX2-1-optimized foregut spheroids express approximately 100-fold more *NKX2-1*. As controls, we included undifferentiated iPSCs and hindgut spheroids, both of which had very low *NKX2-1,* and RNA from fetal lung was used as a positive control. Using an NKX2-1-EGFP reporter cell line, EGFP was not detected at day 7, whereas low-level ubiquitous expression with scattered NKX2-1^HI^ cells could be detected starting on day 10, and expression was localized to specific regions by day 14 (Fig. 1C). Whole mount immunofluorescence (IF) on day 10 spheroids correlated with reporter expression, with individual cells expressing high levels of NKX2-1 protein (Fig. 1D).

**Figure 1:**
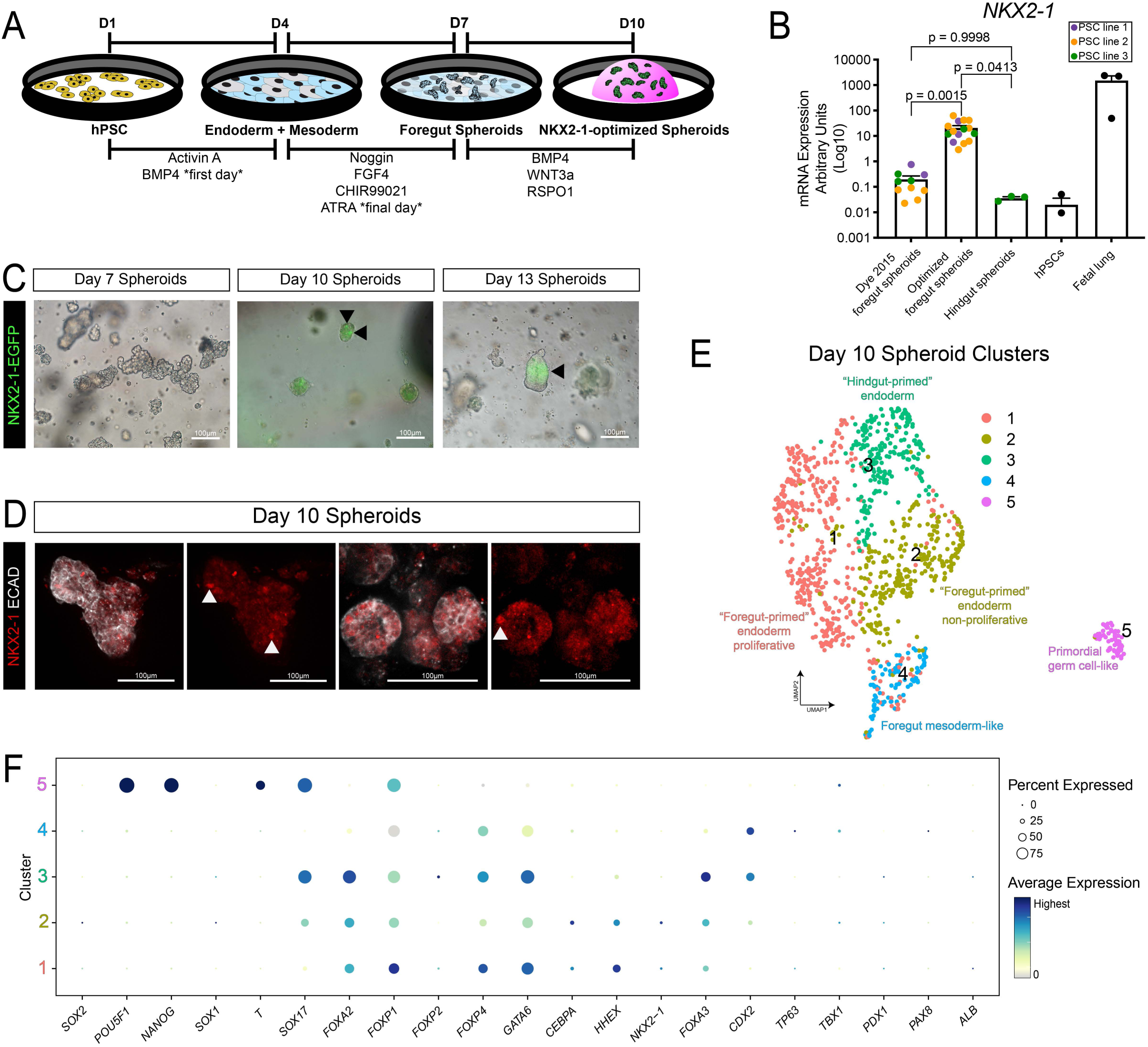
Optimization of Lung Spheroids for NKX2-1 Expression. (A) Schematic displaying the directed differentiation protocol from hPSC to NKX2-1-optimized spheroids. (B) RT-qPCR data comparing *NKX2-1* expression in previously published foregut spheroids (Dye *et al*., 2015) to optimized foregut spheroids and hindgut spheroids (Spence *et al*., 2011). iPSCs and whole fetal lung are also included as references. Each color represents an independent experiment with a unique iPSC line (purple: WTC11, orange: iPSC17 WT 7B2, green: iPSC line 72.3). Each data point of the same color represents a technical replicate from the same iPSC line in one or more independent experiments. Error bars represent standard error of the mean. Statistical tests were performed by ordinary one-way ANOVA followed by Turkey’s multiple comparison test. (C) Representative reporter expression for NXK2-1-EGFP on day 7, 10, or 14 spheroids. (D) Maximum intensity projection of a whole mount immunofluorescence confocal z-series staining for the pan-epithelial marker ECAD and lung epithelial marker NKX2-1. (E) UMAP cluster plot of scRNA-seq data from day 10 spheroids (n=1). Each dot represents a single cell and cells were computationally clustered based on transcriptional similarities. The plot is colored and numbered by cluster. Cell-type labels for each cluster are based on expression of canonical cell-type markers displayed in the dot plot in Fig. 1E or Fig. S1G. (F) Dot plot of cell lineage genes in each cluster of the UMAP plot in Fig. 1D. The dot size represents the percentage of cells expressing the gene in the corresponding cluster, and the dot color indicates log-normalized expression level of the gene.

When day 10 spheroids were analyzed by scRNA-seq, *NKX2-1* was also observed in a subset of cells (Fig. 1E,F), revealing heterogeneity within the foregut spheroids and confirming the findings by EGFP reporter expression. Spheroids included two clusters (Clusters 1 and 2) of lung-fated ventral foregut endoderm-like cells expressing *NKX2-1, FOXA2*, *FOXP1*, *FOXP4* and *HHEX* (Bogue *et al*., 1998; Shu *et al*., 2007; Spence *et al*., 2009; Kearns *et al*., 2013; Davenport *et al*., 2016; Li *et al*., 2016). Clusters and cells were largely negative for other foregut lineage markers including *TP63*, *TBX1*, *PDX1*, *PAX8* and *ALB*; however, there was a clear population of cells that express *FOXA2*, *FOXA3*, *SOX17* and *CDX2* (Cluster 3), which we refer to as “hindgut-primed” endoderm. Additionally, we observed a cluster of *CDX2*^+^/*HAND1*^+^/*ISL1*^+^/*BMP4*^+^/*FOXF1*^+^/*LEF1^+^* (Cluster 4) foregut mesoderm-like cells (Han *et al*., 2020) and a small population of cells (Cluster 5) expressing markers indicative of primordial germ cells, including *POUF1*, *NANOG*, *T*, *SOX17*, *NANOS3* and *TFAP2C* (Davenport *et al*., 2016; Jo *et al*., 2021) (Fig. 1F, S1D). Comparing optimized foregut spheroids to previously published methods (Dye *et al*., 2015; Miller *et al*., 2019) by qRT-PCR, we observed that the hindgut endoderm marker *CDX2* is not statistically different but the posterior foregut endoderm marker *SOX2* (Que *et al*., 2007) is reduced with the optimized method (Fig. S1F). Taken together, our data shows that optimized foregut spheroids have much higher levels of NKX2-1 when compared to prior methods, but they are still heterogeneous with distinct populations of lung-fated cells and hindgut/intestine-fated cells.

### iPSC-derived Bud Tip Progenitors Emerge Over Time

Once NKX2-1^+^ spheroids were formed, we asked how efficiently these spheroids would give rise to BTP-like cells as spheroids expanded into larger organoid structures. Our previous studies have shown that “3 Factor (3F) media” possessing FGF7, CHIR99021 and ATRA expands primary human fetal bud tip progenitors and SOX2^+^/SOX9^+^ iPSC-derived BTP-like cells; however, efficiency and culture over prolonged periods of time were not assessed (Miller *et al*., 2018). After inducing NKX2-1-optimized spheroids on day 10, media was switched to 3F BTP media (Fig. 2A). Spheroids were maintained in 3F for several weeks, where they expanded into complex, branching structures termed “lung progenitor organoids (LPOs),” similar to what we have observed previously (Miller *et al*., 2018) (Fig. 2B). LPOs were passaged every 2 – 4 weeks as either intact organoids with minimum fragmentation (whole-passaged) or were sheared by being drawn through a hypodermic needle or pipette, which is a standard method for passaging primary BTP organoids (Miller *et al*., 2018, 2020; Hein *et al*., 2021). Using an NKX2-1-EGFP reporter iPSC line, to compare retention of lung identity after various passaging methods, only whole-passaged organoids maintained robust EGFP^+^ reporter expression (Fig. 2C). Sheared LPOs recovered well and proliferated faster than whole-passaged LPOs (data not shown); however, fragmenting organoids ultimately led to a loss of NKX2-1-EGFP^+^ cells (Fig. 2C, quantified in Fig. 2F). Based on the maintenance of NKX2-1 expression, we therefore chose to use whole-passaged LPOs for our remaining experiments.

**Figure 2:**
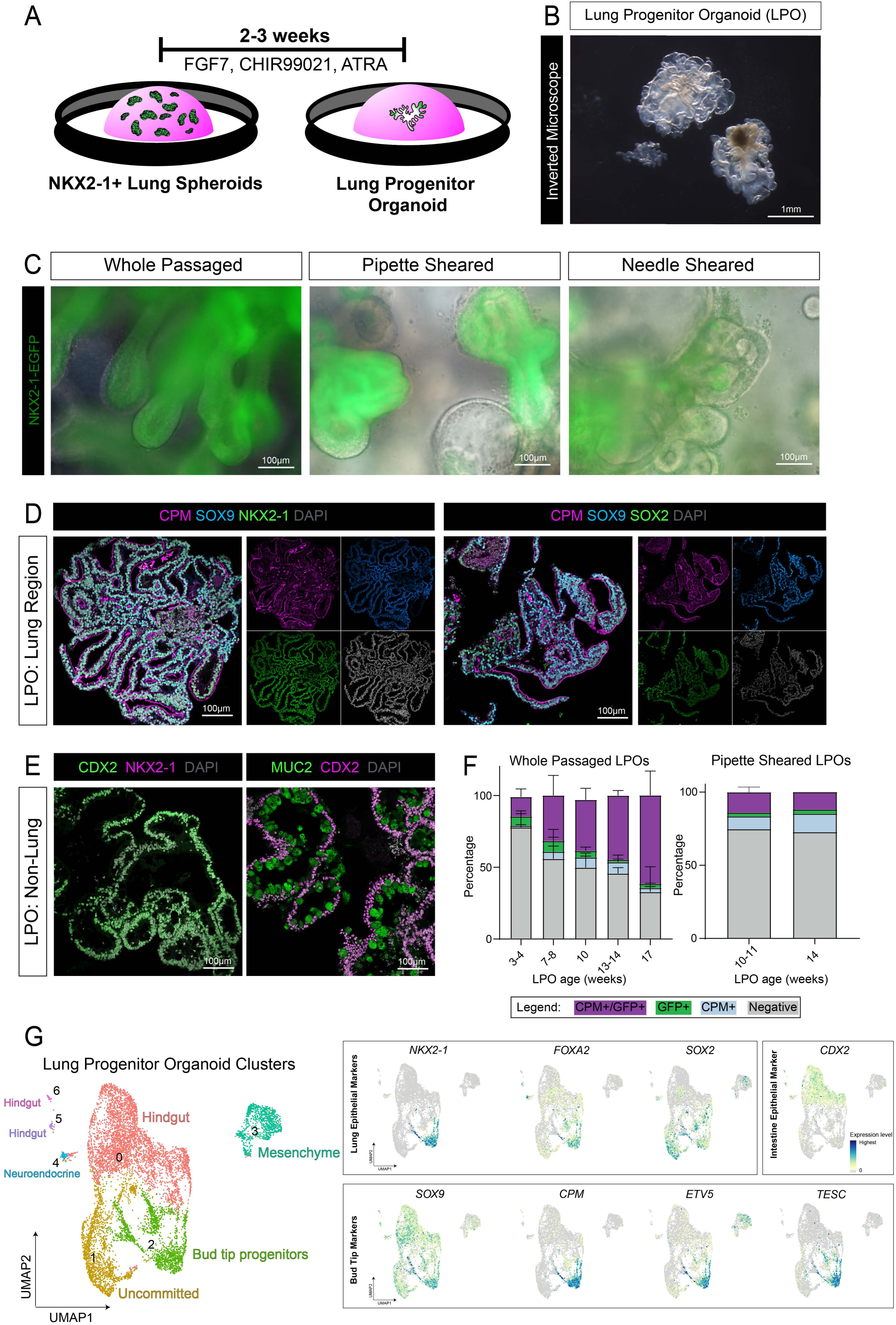
iPSC-derived Bud Tip Progenitors Emerge Over Time in LPOs. (A) Schematic displaying the Lung Progenitor Organoid (LPO) expansion protocol from NKX2-1-optimized spheroids. LPOs form after 2 – 3 weeks in culture. (B) Brightfield image of 6-week LPO on inverted microscope. (C) Representative NKX2-1-EGFP reporter images of 10 – 11-week LPOs, passaged whole or sheared (pipette and needle). (D) Immunofluorescence staining on paraffin sections of the lung-like regions of 12-week LPOs for bud tip progenitor markers CPM and SOX9 and either lung epithelial markers NKX2-1 (left panels) or SOX2 (right panels). (E) Immunofluorescence staining on paraffin sections for intestinal epithelial marker CDX2, intestinal goblet cell marker MUC2, and lung epithelial marker NKX2-1 on non-lung regions of 12-week LPOs. (F) FACS quantification of NKX2-1-EGFP^+^/CPM^+^ cells in LPOs in aggregate time course from the NKX2-1-EGFP reporter cell line. LPOs were sorted using the NKX2-1-EGFP reporter and bud tip progenitor cell surface marker CPM from 3 – 17 weeks in 3F media. Percentages of live cells expressing neither marker (Negative, grey), each separate marker (CPM^+^-only, blue; EGFP^+^-only, green), or dual-expressing cells (CPM^+^/EGFP^+^, purple) are reported as mean ± SEM for 2 – 3 replicates per time point. (G) UMAP cluster plot of scRNA-seq data from LPOs (n=2 biological replicates for 3- and 6-week timepoints, n=1 for 10-week timepoint). Each dot represents a single cell and cells were computationally clustered based on transcriptional similarities. The plot is colored and numbered by cluster. Cell-type labels for each cluster are based on expression of canonical cell-type markers displayed in the heat map in Fig. S3 and the dot plot and feature plots in Fig. 2G and Fig. SE. Feature plots corresponding to the LPO cluster plot and displaying canonical bud tip progenitor markers (*SOX9, CPM, ETV5, TESC*) (Miller *et al*., 2018, 2020; Yamamoto *et al*., 2020), lung epithelial markers (*NKX2-1, FOXA2, SOX2*) and intestine epithelial marker (*CDX2*). The color of each dot in the feature plot indicates log-normalized expression level of the set of bud tip progenitor genes in the represented cell.

We evaluated the presence of BTP-like cells within whole-passaged LPOs by IF of paraffin sections and whole mount (Fig. 2D, S2A). LPOs possessed regions of NKX2-1-expressing cells, which co-expressed the BTP markers CPM, SOX9 and SOX2 (Nikolić *et al*., 2017; Miller *et al*., 2018; Yamamoto *et al*., 2020) (Fig. 2D, S2A). LPOs also contained sub-regions of distinct NKX2-1^-^/CDX2^+^ cells, with many expressing MUC2, indicative of goblet-like cells, and smaller regions of NKX2-1^-^/CDX2^-^ cells (Fig. 2E, S2B).

We quantitatively assessed EGFP^+^/CPM^+^ cells in LPOs grown for 3 – 17 weeks in 3F media using fluorescence activated cell sorting (FACS) (Fig. 2F, S2C,D). We observed an increase in EGFP^+^/CPM^+^ cells in culture over time (Fig. 2F, purple bars). At 3 weeks, 12% of cells expressed CPM and EGFP, in contrast to 17 weeks, where 62% of cells in culture were EGFP^+^/CPM^+^ (Fig. 2F). A small portion (<10%) of cells were singly positive for EGFP (green) or CPM (blue) at any timepoint, and a population of double-negative cells were observed at all times, suggesting that some heterogeneity is maintained in these organoids. We interrogated LPOs derived from two additional non-reporter iPSC lines, using only CPM to quantify BTP-like cells (Fig. S2C). We observed a similar increase of BTP-like cells in culture over time with ∼10% CPM^+^ cells at 3.5 weeks and nearly 50% CPM^+^ cells by 17 weeks (Fig. S2C). Lastly, organoids derived from previously published lung organoid protocols (Dye *et al*., 2015; Miller *et al*., 2019) contained approximately 3-15% CPM^+^ cells at any timepoint examined (Fig. S2C, right panel). The increase in CPM^+^ cells over time suggests that optimized culture conditions promote the emergence and selection of iBTPs.

To further interrogate the heterogeneity and complexity of the LPOs, we performed scRNA-seq on whole-passaged LPOs at 3, 6, and 10 weeks (Fig. 2G, S2E-G, S3). We identified one cluster expressing robust levels of *NKX2-1* and BTP genes (Cluster 2), several clusters expressing hindgut markers (Clusters 0, 5, 6), a mesenchymal cluster (Cluster 3), a neuroendocrine-like cluster (Cluster 4) and a cluster of unknown/uncommitted cells (Cluster 1) (Fig. 2G, S2E-G). The proportion of cells in the BTP cluster (Cluster 2) increased over time while mesenchyme (Cluster 3) was depleted over time and hindgut cells (Clusters 0, 5, 6) were persistent (Fig. S2F). Together this data supports FACS data suggesting that iBTPs continue to expand within LPOs over time and identifies contaminating lineages that persist.

### iPSC-derived Bud Tip Progenitors can be Isolated, Expanded and Maintained Long-term

Although LPOs at every time point contain a significant population of induced BTPs (iBTPs), given the large population and persistence of hindgut lineages, we aimed to isolate NKX2-1-EGFP^+^/CPM^+^ cells to generate higher purity induced bud tip organoid (iBTO) cultures. iBTPs were isolated via FACS with CPM and NKX2-1-EGFP (or CPM only for non-reporter cell lines) and were replated in Matrigel at 8,000 cells/µL in 3F media to form iBTOs (Fig. 3A,B). The NKX2-1-EGFP reporter line showed uniform expression of NKX2-1-EGFP in iBTOs (Fig. 3B) and IF of paraffin sections showed iBTOs contain a near-homogenous population of cells expressing BTP markers CPM, SOX9 and SOX2 (Fig. 3C, S4). Interestingly, the age of LPOs at the time of sorting was a critical determinant of the ability of iBTOs to maintain NKX2-1 and CPM expression. For example, iBTOs generated from 3-week LPOs maintain 4.05% NKX2-1-EGFP^+^/CPM^+^ cells when resorted 7 weeks later. In contrast, iBTOs from >8-week LPOs maintain ∼83% NKX2-1-EGFP^+^/CPM^+^ cells when resorted 8 weeks later (Fig. 3D). This suggests that iBTPs undergo increasing commitment to a bud tip progenitor identity as they are maintained in 3F at the LPO stage.

**Figure 3:**
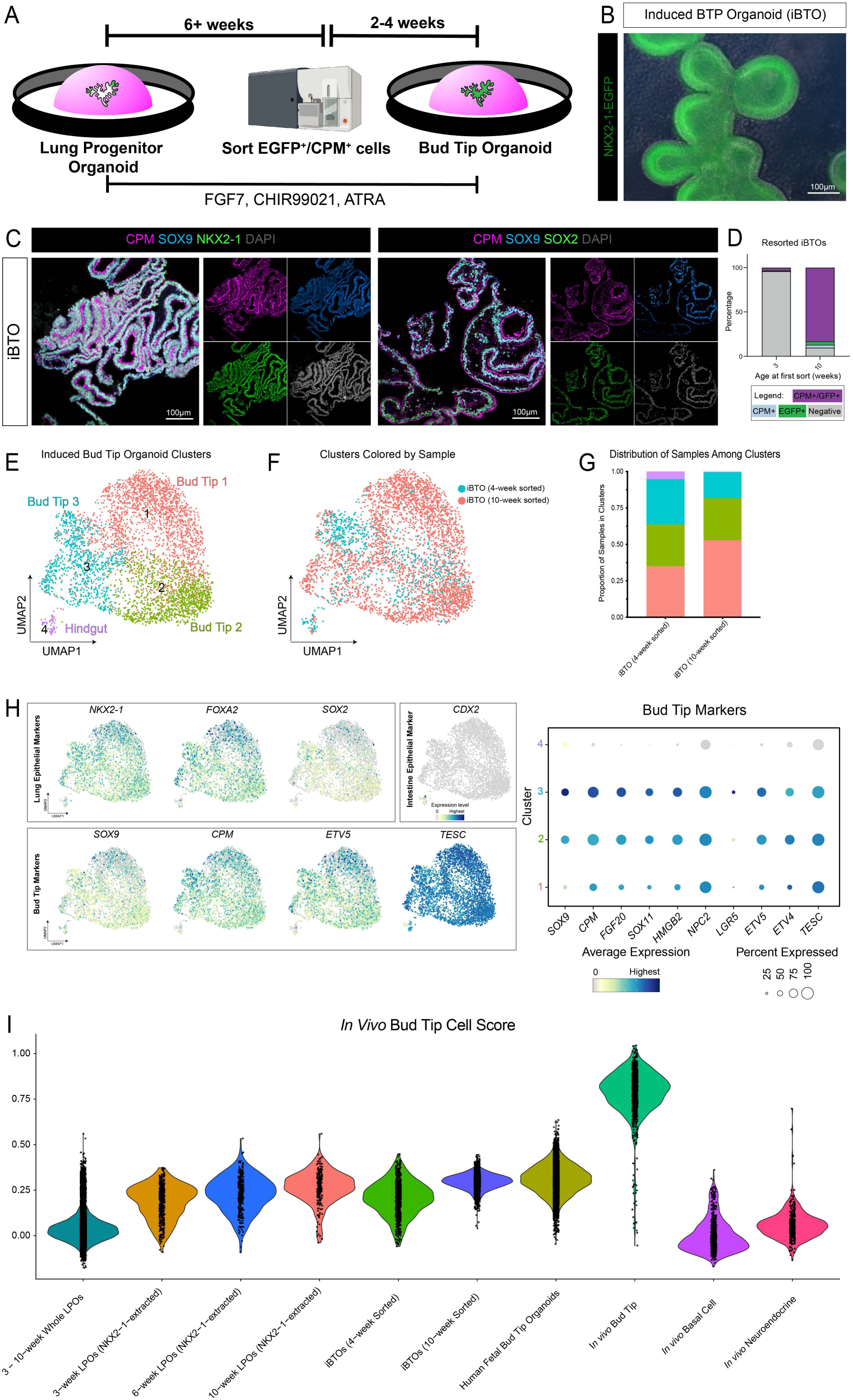
iPSC-derived Bud Tip Progenitors can be Isolated, Expanded Long-term and are Transcriptionally Similar to Human Fetal Bud Tip Progenitor Cultures. (A) Schematic displaying isolation and expansion of induced bud tip progenitor organoids (iBTOs). LPOs are maintained in 3F media and whole passaged for at least 6 weeks then are dissociated for FACS. iBTPs are isolated using CPM^+^ expression +/- NKX2-1EGFP^+^ reporter expression and replated as isolated iBTPs. iBTPs re-form to iBTOs over 2-4 weeks, are maintained in 3F media, and whole passaged. (B) Representative NKX2-1-EGFP reporter image of 3-week iBTO. (C) Immunofluorescence staining on paraffin sections of nearly homogenous 4-week iBTOs for bud tip progenitor markers CPM and SOX9 and either lung epithelial markers NKX2-1 (left panels) or SOX2 (right panels). (D) FACS quantification of NKX2-1-EGFP^+^/CPM^+^ cells in iBTOs from 3-week sorted LPOs or 8 – 17-week sorted LPOs from the NKX2-1-EGFP reporter cell line. iBTOs were analyzed 7 – 8 weeks after iBTP purification from LPOs. Percentages of live cells expressing neither marker (Negative, grey), each separate marker (CPM^+^-only, blue; EGFP^+^-only, green), or dual-expressing cells (CPM^+^/EGFP^+^, purple) are reported as mean ± SEM for 1 – 2 replicates per time point. (E) UMAP cluster plot of scRNA-seq data from iBTOs (n=1 biological replicates for iBTOs from 4- and 10-week LPOs). Each dot represents a single cell and cells were computationally clustered based on transcriptional similarities. The plot is colored and numbered by cluster. Cell-type labels for each cluster are based on expression of canonical cell-type markers displayed in the dot plots and feature plots in Fig. 3H and Fig. S4B. (F) UMAP plot corresponding to the iBTO cluster plot in Fig. 3E. Each dot represents a single cell and dots/cells are colored by the sample from which they came from. (G) Stacked bar graph displaying the proportion of cells from each sample in each cluster of the iBTO cluster plot in Fig. 3E. (H) Feature plots and dot plot corresponding to the iBTO cluster plot in Fig. 3E and displaying canonical bud tip progenitor markers (*SOX9, CPM, ETV5, TESC, FGF20, SOX11, HGMB2, NPC2, LGR5, ETV4*) (Miller *et al*., 2018, 2020; Yamamoto *et al*., 2020), lung epithelial markers (*NKX2-1, FOXA2, SOX2*) and intestine epithelial marker (*CDX2*). The color of each dot in the feature plot indicates log-normalized expression level of the set of bud tip progenitor genes in the represented cell. For the dot plot, the dot size represents the percentage of cells expressing the gene in the corresponding cluster, and the dot color indicates log-normalized expression level of the gene. (I) Violin plot displaying an *in vivo* bud tip progenitor cell score, calculated as the average expression of the top 50 enriched genes in *in vivo* bud tip progenitor cells, for each sample. Samples include whole LPOs (2x3-week LPOs, 2x6-week LPOs, 1x10-week LPOs), NXK2-1-extracted cells from 3-, 6-, and 10-week LPOs (n = 2 for 3- and 6-week LPOs, n = 1 for 10-week LPOs), whole iBTOs (derived from LPOs sorted for NKX2-1^+^/CPM^+^ cells at 4 or 10 weeks, n=1 of each), human fetal-derived bud tip organoids (14-weeks post-conception) (Miller *et al*., 2020), and primary *in vivo* tissue (8.5 – 19-weeks post-conception), including computationally-extracted bud tip, basal, and neuroendocrine cells (Miller *et al*., 2020; Hein *et al*., 2021). The cell scoring list was generated from scRNA-seq data from the *in vivo* bud tip progenitor shown on the plot.

### iPSC-derived Bud Tip Progenitors are Transcriptionally Similar to Human Fetal Bud Tip Progenitor Cultures

To further interrogate iBTO cellular composition, and to directly compare cells within iBTOs to primary human fetal bud tip progenitors and organoids derived from fetal bud tips (Miller *et al*., 2018), we performed scRNA-seq on iBTOs derived from 4- and 10-week LPOs. iBTOs were expanded for 4 weeks in culture before scRNA-seq was carried out. Integrated analysis of both data sets resulted in 4 epithelial clusters, (Fig. 3E, S4B). Cluster 4 expressed hindgut/intestinal markers (i.e. *CDX2*), which could also be identified by immunofluorescence (Fig. S4A) and represented a small fraction of cells that are predominantly derived from the 4-week sample (Fig. 3G). The remaining 3 clusters (Clusters 1, 2, 3) express *NKX2-1* as well as BTP markers (*SOX9, CPM, ETV5, TESC*, *FGF20*, *SOX11*, *HMGB2*, *NPC2*, *LGR5*, *ETV4*) (Miller *et al*., 2018, 2020; Hein *et al*., 2021); however, they also expressed proliferation genes at varying levels, suggesting there is some heterogeneity within iBTPs that is likely driven by the expression of proliferation genes (Fig. 3H).

To benchmark transcriptional similarities iBTPs to *in vivo* BTPs, we re-analyzed published scRNA-seq data from the human fetal lung (Miller *et al*., 2020; Hein *et al*., 2021) and identified a panel of the top 93 enriched genes expressed in *in vivo* BTPs relative to all other epithelial cell types in the developing lung (See ‘Methods’). We used this gene list as a reference to assign an “*in vivo* bud tip progenitor cell score” to each individual cell from the LPOs and iBTOs using published computational methods (Holloway *et al*., 2020; Hein *et al*., 2021). For the purposes of this comparison, both whole LPOs and extracted *NKX2-1*^+^ cells from the LPO data sets were included in the analysis. As additional control comparisons, we analyzed published data and included scores for BTOs derived from fetal lung, *in vivo* basal cells and *in vivo* neuroendocrine cells (Miller *et al*., 2018, 2020; Hein *et al*., 2021) (Fig. 3I). As expected, *in vivo* BTPs had the highest score (median score = 0.80) while *in vivo* basal cells, *in vivo* neuroendocrine cells and whole LPOs had the lowest scores (−0.01, 0.05, 0.03 respectively). A noteworthy observation is that human fetal BTOs had a mean BTP score of 0.31, which was much lower than the *in vivo* BTP score of 0.80, indicating that the *in vitro* culture conditions significantly influence gene expression, as has been observed previously (Miller *et al*., 2020). Relative to fetal-derived BTOs, both *NKX2-1*^+^ cells from LPOs and iBTOs show high transcriptional similarity; their scores improve with longer time in culture at the LPO stage (0.21 to 0.27 for *NKX2-1^+^* cells from LPOs; 0.21 to 0.30 for iBTOs) (Fig. 3I). To provide additional confidence in this comparison method, we generated a similar *in vivo* basal cell score and *in vivo* neuroendocrine cell score, and observed that the expected populations (i.e. *in vivo* basal cells and *in vivo* neuroendocrine cells, respectively) scored the highest (Fig. S4C,D). Taken together, our data supports the conclusion that iBTOs represent a stable bud tip progenitor cell that can be maintained *in vitro* and that share a high degree of transcriptional similarity to fetal bud tip progenitors.

### iBTOs Can Give Rise to Airway or Alveolar Fates

Given that BTPs in the developing lung give rise to alveolar and airway fates, we hypothesized that iBTOs could be guided into airway and alveolar lineages. To test this possibility, we used methods previously developed to efficiently induce lung progenitors into airway (Miller *et al*., 2020) and alveolar lineages (Jacob *et al*., 2017) (Fig. 4A). Airway induction involved 3 days of dual-SMAD activation (DSA) followed by 18 days of dual-SMAD inhibition (DSI) and resulted in condensed structures that maintained NKX2-1-EGFP expression and expressed mCherry driven by the TP63 promoter (TP63-mCherry) (Fig. 4B, left column). By immunofluorescence, we observed that TP63 expression was highly induced after 3 days of DSA, as expected, and after 18 days of DSI, TP63^+^ cells organized around the outside of the organoids (Fig. 4C, S4B).

**Figure 4:**
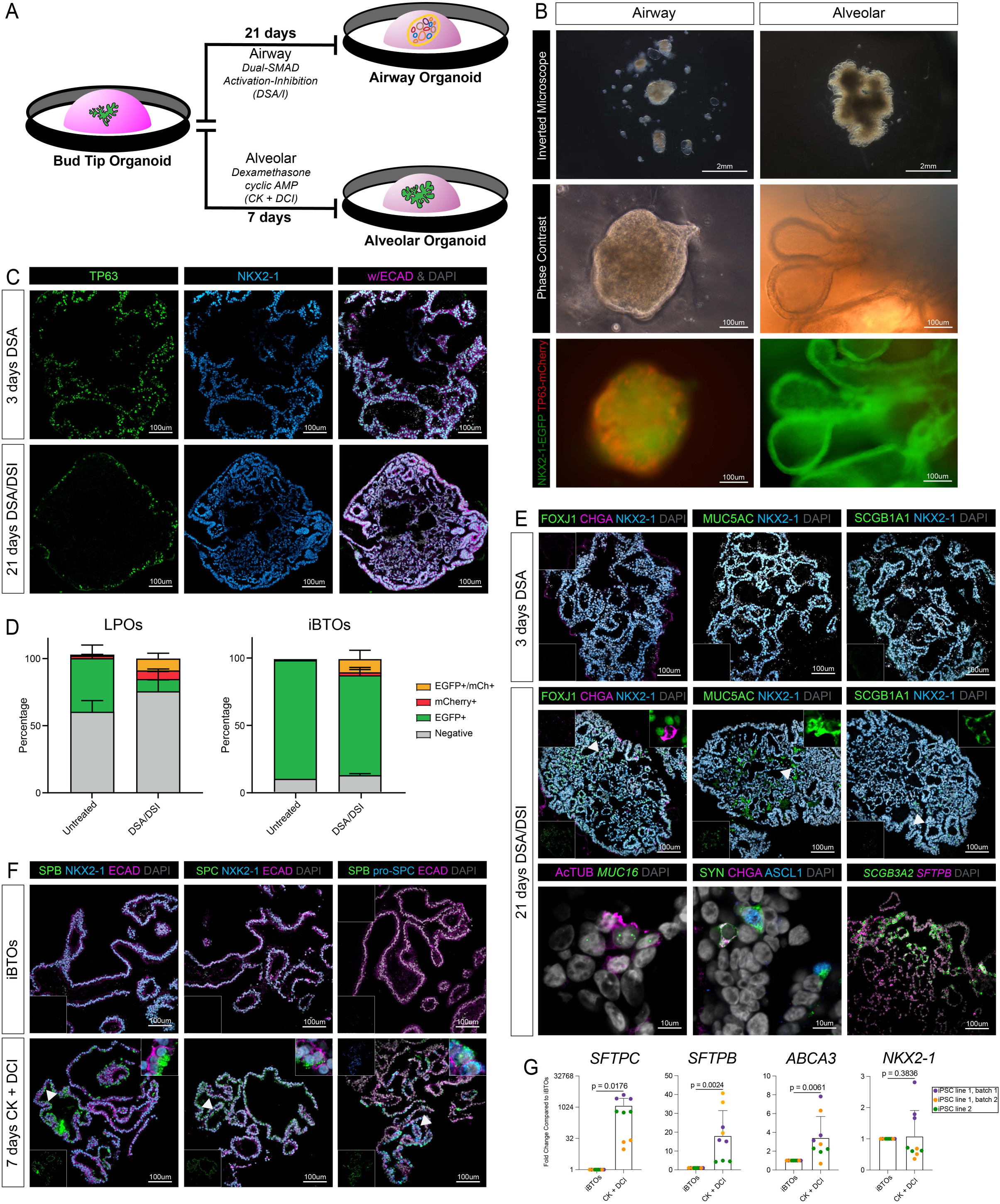
iBTOs are Competent for Proximal Airway and Distal Alveolar Differentiation. A) Schematic displaying the airway (Miller *et al*., 2020) and alveolar (Jacob *et al*., 2017) induction protocol from iBTOs. B) Representative brightfield, phase contrast, and reporter expression for NKX2-1-EGFP and TP63-mCherry images of iBTOs undergone the airway or alveolar induction protocol shown in Fig. 4A. Phase contrast and reporter images are from the same field. C) Immunofluorescence staining on paraffin sections for the airway progenitor marker TP63 and lung epithelial marker NKX2-1 on iBTOs undergone 3 days of dual-SMAD activation (DSA) or 3 days DSA followed by 18 days of dual-SMAD inactivation (DSI) in the airway induction protocol. D) FACS quantification of NKX2-1-EGFP^+^ and mCherry-TP63^+^ cells from LPOs and iBTOs from the NKX2-1-EGFP/TP63-mCherry reporter cell line, either in 3F media (untreated) or after 21 days of the airway differentiation protocol (DSA/DSI treatment). Percentages of live cells expressing neither marker, each separate reporter, or dual-expressing cells are reported as mean ± SEM for 2 – 3 replicates per time point. E) Fluorescence *in situ* hybridization and/or immunofluorescence staining on paraffin sections for differentiated airway epithelial markers (multiciliated: FOXJ1/AcTUB/*MUC16*, neuroendocrine: CHGA, SYN, ASCL1, goblet: MUC5AC, secretory: SCGB1A1, *SCGB3A2*, *SPB*) and lung epithelial marker NKX2-1 on iBTOs undergone 3 DSA or 3 days DSA followed by 18 days of DSI of the airway induction protocol. Insets on the top right are zoomed in on the area denoted by the arrows. Insets in the bottom left or top left corners are single channel images. F) Immunofluorescence staining on paraffin sections for differentiated type II alveolar epithelial markers SPB, pro-SPC, and SPC and lung epithelial marker NKX2-1 or general epithelial marker ECAD on iBTOs and iBTOs undergone the alveolar differentiation protocol (7 days CK + DCI). Insets on the top right are zoomed in on the area denoted by the arrows. Insets in the bottom left or top left corners are single channel images. G) RT-qPCR data comparing expression of alveolar markers *SFTPC, SFTPB,* and *ABCA3* and lung epithelial marker *NKX2-1* in untreated iBTOs (in 3F media) or after the alveolar differentiation protocol (7 days CK + DCI). Each color represents an independent experiment with a unique iPSC line or independent experiment (purple/orange: iPSC17 WT 7B2 – LPOs sorted by CPM and NKX2-1-EGFP, green: iPSC line 72.3 – LPOs sorted by CPM only). Error bars represent standard error of the mean. Statistical tests were performed by Student’s t-test (paired, one-tailed).

We performed airway induction on unsorted LPOs as well, hypothesizing that airway differentiation may select for expansion of lung lineages within the LPO, while suppressing non-lung lineages.TP63-mCherry^+^ cells were quantified using flow cytometry on both LPOs and iBTOs following the 21-day DSA/DSI protocol or untreated controls (Fig. 4D). Untreated LPOs contained 0.36% NKX2-1-EGFP^+^/TP63-mCherry^+^ cells while untreated iBTOs contained 0.76%. Only upon treatment with the 21-day DSA/DSI protocol were significant numbers of EGFP^+^/mCherry^+^ cells detected (9% in LPOs, 16% in iBTOs) (Fig. 4D, S4B). DSA/DSI-treated iBTOs also maintained a large portion (70%) of NKX2-1-EGFP^+^/TP63-mCherry^-^ cells, which represented the spectrum of differentiated airway epithelial cell types identified by immunofluorescence, including multiciliated, neuroendocrine, goblet and secretory cells (Fig. 4D,E). Airway organoids contained AcTUB^+^/*MUC16^+^* ciliated cells (Carraro *et al*., 2021) (Fig. 4E), and SYN^+^ cells were double positive for ASCL1 (early neuroendocrine marker) or CHGA (late neuroendocrine marker) (Fig. 4E). Treated LPOs contained a small proportion of cells only expressing TP63-mCherry (6%) (Fig. 4D, S4B) and majority were negative for both markers (76%), suggesting non-lung lineages continue to expand in these conditions

Alveolar induction consisted of 7 days of treatment with CK + DCI as has been previously described with the exception of an alternate composition of the base media (see ‘Methods’) (Jacob *et al*., 2017). Treatment of iBTOs with CK + DCI resulted in expanded budded structures that maintained NXK2.1-EGFP expression (Fig. 4B, right column). Alveolar type II markers *SFTPB*, *SFTPC* and *ABCA3* were low to undetectable in iBTOs but were highly expressed upon treatment with CK + DCI as determined by qRT-PCR and co-expressed within the same cells as shown by immunofluorescence for (pro-)SFTPC, SFTPC, SFTPB and NKX2-1 (Fig. 4F,G). The lung-specific epithelial marker NKX2-1 was robust in both airway and alveolar organoids while expression of the intestinal epithelial marker CDX2 was not regularly detected (Fig. 4D-G, S4A). Taken together, differentiation of iBTOs into airway and alveolar cell types with minimal contaminating non-lung cell types supports that iBTOs have similar developmental potential to *in vivo* bud tip progenitors.

## DISCUSSION

With the current work, we demonstrate an optimized *in vitro* model system for differentiating iPSCs into an NKX2-1-expressing bud tip progenitor lineage, which we show can give rise to both airway and alveolar cell types. Several groups, including ours (Dye *et al*., 2015, 2016; Miller *et al*., 2018, 2019), have previously characterized protocols that yield lung-like cell types from iPSCs (Wang *et al*., 2007; Mou *et al*., 2012; Gotoh *et al*., 2014; Huang *et al*., 2014; Konishi *et al*., 2016; Abler *et al*., 2017; Chen *et al*., 2017; Hawkins *et al*., 2017; McCauley *et al*., 2017; Tamò *et al*., 2018; de Carvalho *et al*., 2019; Jacob *et al*., 2019; Yamamoto *et al*., 2020; Leibel *et al*., 2020). While many of these protocols capture a transient bud tip progenitor-like stage, they focus primarily on deriving more mature lung epithelial lineages, and the induction and maintenance of NKX2-1^+^ bud tip progenitor-like cells from iPSCs has not been previously reported. Furthermore, benchmarking of induced cells using scRNA-seq in order to compare cell types to a human reference atlas has only recently become commonplace (Holloway *et al*., 2020; Hor *et al*., 2020; Yu *et al*., 2020; Hawkins *et al*., 2021). The method reported here expands upon an improved understanding of critical signaling events that regulate bud tip progenitor maintenance in the human lung to better mirror this process in the dish (Conway *et al*., 2020).

Although we demonstrate that optimizing NKX2-1 expression leads to a 100-fold induction when compared to previously reported methods (Dye *et al*., 2015; Miller *et al*., 2019), scRNA-seq and whole mount immunofluorescence analysis of day 10 spheroids revealed that low levels of NKX2-1 is expressed in many cells, while a small number of cells express high levels of NKX2-1, highlighting the heterogeneity that still exists at this early timepoint. We also observed non-lung lineages that co-emerge in spheroids, possibly due to the fact that the same signaling pathways play a role during lineage choice for multiple lineages, and due to the inherent plasticity of newly committed cells. We observed that non-lung lineages, and particularly gut lineages, persisted over time in culture. Based on this observation, there is an opportunity to further refine the signaling pathways that are manipulated to expand and maintain NKX2-1^+^ lung lineages; however, this may prove challenging given the redundant use of some pathways (i.e. WNT) to maintain stem/progenitor cells from both lineages (Pinto and Clevers, 2005; Kapoor, Li and Leiter, 2007; Volckaert and De Langhe, 2015; Chin *et al*., 2016; Nikolić and Rawlins, 2017; Yan *et al*., 2017; Miller *et al*., 2018; Rabata *et al*., 2020; Aros, Pantoja and Gomperts, 2021). We observed that hindgut lineages expand after passaging organoids using the standard approach of mechanically shearing them into small fragments. It is currently unclear if this phenomenon occurs because shearing increases selection for highly proliferative CDX2^+^ cells or if the mechanical stress results in higher levels of apoptosis in one lineage (i.e. lung) compared to the other. Thus, while we have optimized the current methods, to generate a high proportion of NKX2-1^+^ cells, LPO cultures still exhibit high plasticity that can be triggered through perturbations such as organoid dissociation.

One important observation from our current study is that iBTPs exhibit increased lineage commitment with longer time in culture. Isolating NKX2-1-EGFP^+^/CPM^+^ cells prior to 3 weeks resulted in cultures replete with gut cells. While follow-up experiments are required to determine the origin of contaminating gut cells, the fact that we still see these cells in iBTOs suggests a fate switch from a lung bud tip progenitor to a hindgut identity. On the other hand, purifying cells at 6 – 10 weeks reliably led to the establishment of NKX2-1-EGFP^+^/CPM^+^ iBTOs that expand and maintain their fate. We attempted airway differentiation on unsorted LPOs, hypothesizing that the process of differentiation itself may select against gut lineages and expand only lung lineages; unfortunately, non-lung lineages persisted. Given this result, sorting remains an important step in establishing iBTO cultures that can be differentiated into airway or alveolar cell types in the absence of non-lung cell types. Finally, the experiments carried out here relied heavily on an NKX2-1-EGFP reporter; however, our results suggest that this should not limit studies with non-reporter iPSC lines, since we also observed that sorting with CPM alone highly enriched for iBTPs. As we detected some NKX2-1-EGFP^-^/CPM^+^, identifying additional bud tip progenitor cell surface markers to further improve purity of cultures may be valuable for enhancing purity in non-reporter iPSC cultures.

The utilization of emerging technologies such as scRNA-seq has provided critical insights into the heterogeneity and complexity of human tissues that were once difficult to study (Miller *et al*., 2020; Yu *et al*., 2020). New information from human tissue has provided both an atlas against which *in vitro* model systems can be benchmarked and a roadmap that can be used to infer transcriptional and signaling mechanisms that control cellular transitions. Here, we used scRNA-seq to benchmark *in vitro* cultures at several stages during differentiation to catalog the induction, emergence, and maintenance of lung-fated cells as they acquire a bud tip progenitor fate. One interesting observation from the benchmarking carried out in this study is a significant shift in the transcriptome when comparing primary *in vivo* tissue to *in vitro* grown primary organoids, even when the source of the cells is the same. For example, here, we compared iBTOs to BTOs derived from fetal lung and to *in vivo* fetal bud tip progenitors. We observed a significant shift when comparing fetal lung bud tip progenitors to primary BTOs, indicating that the *in vitro* environment significantly changes the transcriptome of the cell, as has been recently reported (Miller *et al*., 2020; Alysandratos *et al*., 2022). With this caveat in mind, iBTOs shared a nearly identical transcriptome to primary BTOs, and shared a high degree of similarity to *in vivo* primary bud tip progenitors. The cell scoring metric used to compare similarity across samples also suggested that iBTOs increased in transcriptional similarity to *in vivo* bud tip progenitors as they spent more time in culture, supporting the idea that iBTPs undergo continued differentiation towards a bud tip progenitor identity as they are maintained in culture.

Functionally, iBTOs also behave like bud tip progenitors. Using methods to differentiate lung progenitor cells into airway or alveolar cell types (Jacob *et al*., 2017; Miller *et al*., 2020) (Miller *et. al.*, 2020, Jacob *et al*., 2017), we demonstrated that iBTOs robustly differentiate into airway and alveolar cell types. Taken together, the current work describes a robust iPSC-derived bud tip progenitor model to study human lung epithelial development and differentiation and uses scRNA-seq to benchmark both on- and off-target cell types present in the cultures. Practically, this model presents a novel opportunity to expand the downstream efforts of lung organoid studies, where iBTOs can be shared to reduce the expertise needed for lung cell generation and ultimately facilitate faster turn-around through iBTO banking. Overall, this study enhances the utility of iPSC-derived lung organoids to interrogate lung development and to study disease and regeneration.

## MATERIALS AND METHODS

### Tissue processing, Staining, and Quantification

All sectioned fluorescent images were taken using a Nikon A1 confocal microscope, an Olympus IX83 inverted fluorescence microscope or an Olympus IX71 inverted fluorescence microscope. Whole mount fluorescent images were taken on a Nikon X1 Yokogawa Spinning Disk Microscope. Acquisition parameters were kept consistent for images in the same experiment and all post-image processing was performed equally on all images in the same experiment. Images were assembled in Adobe Photoshop CC 2022.

#### Tissue Processing

Tissue was immediately fixed in 10% Neutral Buffered Formalin (NBF) for 24h at room temperature on a rocker. Tissue was then washed 3x in UltraPure DNase/RNase-Free Distilled Water (Thermo Fisher, Cat#10977015) for 15 minutes each and then dehydrated in an alcohol series of concentrations dehydrated in UltraPure DNase/RNase-Free Distilled Water for 1h per solution: 25% Methanol, 50% Methanol, 75% Methanol, 100% Methanol, 100% Ethanol, 70% Ethanol. Dehydrated tissue was then processed into paraffin blocks in an automated tissue processor (Leica ASP300) with 1h solution changes. 4 (FISH) or 7 (IF) µm-thick sections were cut from paraffin blocks onto charged glass slides. For FISH, microtome and slides were sprayed with RNase Away (Thermo Fisher, Cat#700511) prior to sectioning (within one week of performing FISH). Slides were baked for 1h in 60°C dry oven (within 24h of performing FISH). Slides were stored at room temperature in a slide box containing a silica desiccator packet and the seams sealed with paraffin.

#### Immunofluorescence (IF) Protein Staining on Paraffin Sections

Tissue slides were rehydrated in Histo-Clear II (National Diagnostics, Cat#HS-202) 2x for 5 minutes each, followed by serial rinses through the following solutions 2x for 2 – 3 minutes each: 100% EtOH, 95% EtOH, 70% EtOH, 30%EtOH, and finally in double-distilled water (ddH2O) 2x for 5 minutes each. Antigen retrieval was performed using 1X Sodium Citrate Buffer (100mM trisodium citrate (Sigma, Cat#S1804), 0.5% Tween 20 (Thermo Fisher, Cat#BP337), pH 6.0), steaming the slides for 20 minutes, followed by cooling and washing quickly 2x in ddH2O and 2x in 1X PBS. Slides were incubated in a humidified chamber at RT for 1h with blocking solution (5% normal donkey serum (Sigma, Cat#D9663) in PBS with 0.1% Tween 20). Slides were then incubated in primary antibody diluted in blocking solution at 4°C overnight in a humidified chamber. Next, slides were washed 3x in 1X PBS for 5 minutes each and incubated with secondary antibody with DAPI (1ug/mL) diluted in blocking solution for 1h at RT in a humidified chamber. Slides were then washed 3x in 1X PBS for 5 minutes each and mounted with ProLong Gold (Thermo Fisher, Cat#P36930) and imaged within 2 weeks. Stained slides were stored in the dark at 4°C. Secondary antibodies, raised in donkey, were purchased from Jackson Immuno and used at a dilution of 1:500.

#### Fluorescence in situ hybridization (FISH)

The FISH protocol was performed according to the manufacturer’s instructions (ACDbio, RNAscope multiplex fluorescent manual) with a 5-minute protease treatment and 15-minute antigen retrieval. For IFco-staining with antibodies, the last step of the FISH protocol was skipped and instead the slides were washed 1x in PBS followed by the IF protocol above.

#### Whole Mount IF Protein Staining

All tips and tubes were coated with 1% BSA in PBS to prevent tissue sticking. 3D cultures (spheroids, LPOs) were dislodged from Matrigel using a P1000 cut tip and transferred to 1.5mL Eppendorf tube. 500µL of Cell Recovery Solution (Corning, Cat#354253) was added to the tube and placed on a rocker at 4°C for 45 minutes to completely dissolve Matrigel. Tube was spun at 100g for 5 minutes, and solution and remaining Matrigel was then removed. Tissue was fixed in 10% NBF overnight at RT on a rocker. Tissue then was washed 3x for 2hrs with 1mL room temperature Organoid Wash Buffer (OWB) (0.1% Triton, 0.2% BSA in 1X PBS), at RT on a rocker. Wash times vary (30 minutes – 2 hours) depending on tissue size. 1mL CUBIC-L (TCI Chemicals Cat#T3740) was added to the tube and placed on a rocker for 24hrs at 37°C. Tissue was then permeabilized for 24hr at 4°C on rocker with 1mL permeabilization solution (5% Normal Donkey Serum, 0.5% Triton in 1X PBS). After 24hr, permeabilization solution was removed and 500µL primary antibody (diluted in OWB) was added overnight at 4°C on a rocker. The next day, tissue was washed 3x with 1mL of OWB, 2hr each at RT. 500µL of secondary antibody (diluted in OWB) was added and incubated overnight at 4°C, wrapped in foil. Tissue was washed again 3x with 1mL OWB at RT, first wash for 2hrs, then 30 mins for the remaining washes. Samples were transferred to imaging plate (ThermoFisher Cat#12-566-70) and then cleared and mounted with 50µL CUBIC-R (just enough to cover tissue) (TCI Chemicals Cat#T3741).

IF and FISH stains were repeated on at least 3 independent differentiations and representative images are show.

### Cell Lines & Culture Conditions

#### hPSC Lines and Culture Conditions

##### hPSC Lines

LPOs were generated from three human iPSC lines: WTC11 (RRID: CVCL_Y803) was obtained from Bruce Conklin at the University of California San Francisco (Kreitzer *et al*., 2013), human iPSC line 72.3 was obtained from Cincinnati Children’s Hospital Medical Center (McCracken *et al*., 2014) and iPSC17 WT 7B2, expressing NKX2-1-EGFP and TP63-mCherry, was obtained from the Cystic Fibrosis Foundation (Crane *et al*., 2015). In addition to iPSC lines, hESC lines were used for definitive endoderm and spheroid optimizations: hESC line H9 (NIH registry no. 0062) was obtained from the WiCell Research Institute and hESC line RUES2-GLR (NIH registry no. 0013), expressing SOX2-mCitrine, BRA-mCerulean and SOX17-tdTomato, was obtained from The Rockefeller University (Martyn *et al*., 2018). The University of Michigan Human Pluripotent Stem Cell Research Oversight Committee approved all experiments using human embryonic stem cell (hESC) and induced pluripotent stem cell (iPSC) lines. Stem cell lines were routinely karyotyped, and all cell lines are routinely monitored for mycoplasma infection monthly using the MycoAlert Mycoplasma Detection Kit (Lonza, Cat#LT08-118). Stem cells were maintained as previously described (Spence *et al*., 2011) and grown in mTeSR Plus media (StemCell Technologies, Cat#100-0276).

##### NKX2-1-optimized Spheroid Differentiation Protocol

Generation of definitive endoderm (DE) from hPSCs (differentiation days 1 – 3) was carried out as previously described with slight modifications (D’Amour *et al*., 2006; McCracken *et al*., 2011; Spence *et al*., 2011; Dye *et al*., 2015). Briefly, 100ng/mL ActivinA (R&D Systems, Cat#338-AC) was added in RPMI 1640 media (Thermo Fisher, Cat#21875034) with increasing concentrations of HyClone defined FBS (dFBS) (Fisher Scientific, Cat#SH3007002) on subsequent days (0% day 1, 0.2% day 2, 2% day 3). 50ng/mL BMP4 (R&D Systems, Cat#314-BP) was added on day 1. Following DE specification, anterior foregut spheroids were generated (differentiation days 4 – 6) by a 3-day treatment with 500ng/mL FGF4, 200ng/mL NOGGIN (R&D Systems, Cat#6057-NG), 2uM CHIR99021 (APExBIO, Cat#A3011) and 2uM all trans retinoic acid (Sigma, Cat#R2625) on the final day. Self-organizing 3D spheroids that had detached from the tissue culture dish were collected with a P200 pipette and were transferred into Matrigel (Corning, Cat#354234) as previously described (McCracken *et al*., 2011; Miller *et al*., 2019). After the Matrigel had solidified, encapsulating spheroids, they were cultured in media that contained 250ng/mL WNT3a (R&D Systems, Cat#5036-WN), 500ng/mL RSPO1 and 10ng/mL BMP4 for 3 days to induce NKX2-1^+^ cells (differentiation days 7 – 9). Following day DE specification, RPMI 1640 media + 2% dFBS was used on all days to dilute growth factors. Fresh media and growth factors were added each day.

##### Growth and Maintenance of LPOs

NKX2-1-optimized spheroids were transferred to “3 Factor” (3F) bud tip maintenance media as previously described (Miller *et al*., 2018, 2019), including 50nM all trans retinoic acid (Sigma, Cat#R2625), 10ng/mL FGF7 (R&D Systems, Cat#251-KG) and 3µM CHIR99021(APExBIO, Cat#A3011) in serum-free basal media. Serum-free basal media consists of DMEM/F12 containing HEPES and L-Glutamine (Corning, Cat#10-092-CV), 100U/mL penicillin-streptomycin (Thermo Fisher, Cat#15140122), 1x B-27 supplement (Thermo Fisher, Cat#17504044), 1x N-2 supplement (Thermo Fisher, Cat#17502048), 0.05% BSA (Sigma, Cat#A9647), 50µg/mL L-ascorbic acid (Sigma, Cat#A4544) and 0.4µM 1-Thioglycerol (Sigma, Cat#M1753). LPOs were grown for 3 weeks, then whole passaged or needle or pipette sheared every 2 – 4 weeks. Whole passaging was achieved by collecting LPOs into an Eppendorf tube and gently releasing them from Matrigel by a P1000 cut pipette tip. LPOs were spun in a microcentrifuge tube, residual media and Matrigel was removed, then LPOs were re-suspended in Matrigel (Corning, Cat#354234) with a P1000 cut pipette tip. ∼35µL droplets of Matrigel were placed into the center of wells of a 24-well tissue culture plate (Thermo Fisher, Cat#12565163), and the plate was inverted and placed in an incubator at 37°C for 20 minutes. For LPOs passaged by needle or pipette shearing, LPOs were passed through a 27-gauge needle or P200 pipette, respectively, and embedded in fresh Matrigel as previously described (Miller *et al*., 2018, 2019, 2020). LPOs were fed with 3F media every 2 – 4 days.

##### Growth and Maintenance of iBTOs

iBTs in LPOs grown in 3F media for 3 – 17 weeks were isolated by FACS (see below). After collection, iBTs were centrifuged at 300g for 5 minutes at 4°C and media was removed. Matrigel (Corning, Cat#354234) was added to the cells at a concentration of 8,000 cells/µL Matrigel and 3 – 20µL droplets of Matrigel were placed into the center of wells of a 24-well tissue culture plate (Thermo Fisher, Cat#12565163). The plate was inverted and placed in an incubator at 37°C for 20 minutes. iBTPs were fed every 2 – 4 days with 3F media and allowed to reform organoids. iBTOs were whole passaged as described above every 2 – 4 weeks.

##### Airway and Alveolar Differentiations

Airway differentiation was carried out as previously described (Miller *et al*., 2020). Briefly, iBTOs were exposed to dual-SMAD activation via 100ng/mL BMP4 (R&D Systems, Cat#314-BP-050) and 100ng/mL TGFB1 (R&D Systems, Cat#240-B-002) in 3F media for 3 days. On the fourth day, iBTOs were exposed to dual-SMAD inactivation via 1µM A8301 (APExBIO, Cat#3133), 100ng/mL NOGGIN (R&D Systems, Cat#6057), 10µM Y-27632 (APExBIO, Cat#B1293) and 500ng/mL FGF10 in serum-free basal media for 18 days (media changed every 3 – 4 days) with whole passaging as necessary.

Alveolar differentiation was carried out as previously described (Jacob *et al*., 2017). Briefly, iBTOs were transitioned to alveolar differentiation media for 7 days (media changed on days 3 and 6). Alveolar differentiation media consists of a modified serum-free basal media (SFB-VA) with DMEM/F12 containing HEPES and L-Glutamine (Corning, Cat#10-092-CV) supplemented with 1X N-2 supplement (Thermo Fisher, Cat#17502048), 1X B-27 supplement without vitamin A (Thermo Fisher, Cat#12587010), 0.05% BSA (Sigma, Cat#A9647), 100U/mL Penicillin-Streptomycin (Thermo Fisher, Cat#15140122), 50µg/mL L-Ascorbic Acid (Sigma, Cat#A9647) and 0.4µM 1-Thioglycerol (Sigma, Cat#M6145). On the first day of alveolar differentiation, SFB-VA is supplemented with 10ng/mL FGF7 (R&D Systems, Cat#251-KG/CF), 3µM CHIR99021 (APExBIO, Cat#A3011), 100µM 3-Isobutyl-1-methylxanthine (Sigma, Cat#I5879), 100µM 8-bromoadenosine 3’, 5’ -cyclic monophosphate sodium salt (Sigma, Cat#B7780) and 50nM Dexamethasone (Sigma, Cat#D4802).

### Culture Media, Growth Factors, Small Molecules

#### Tissue Prep for scRNA-seq

All tubes and pipette tips were pre-washed in 1X HBSS with 1% BSA to prevent cell adhesion to the plastic. 3D cultures (spheroids, LPOs, iBTOs) were removed from Matrigel using a P1000 pipette tip and vigorously pipetted in a 1.5mL Eppendorf tube to remove as much Matrigel as possible. Tissue was centrifuged at 300g for 3 minutes at 4°C, then excess media and Matrigel was removed. Tissue was digested to single cells using 0.5mL TrypLE (Invitrogen, Cat#12605010) and incubated at 37°C for 30 minutes, adding mechanical digestion with pipette every 10 minutes. After 30 minutes, trypsinization was quenched with 1X HBSS. Cells were passed through a 40µm filter (Bel-Art Flowmi, Cat#136800040), and centrifuged at 300g for 3 minutes at 4°C. Cells were resuspended in 1mL 1X HBSS and counted using a hemocytometer, centrifuged at 300g for 3 minutes at 4°C and resuspended to a final concentration of 1,100 cells/µL. If samples were planned for combined submission, cells would be cryopreserved in XX media and stored in liquid N2. Cells were thawed, washed and sent for sequencing. Approximately 100,000 cells were put on ice and single cell libraries were immediately prepared at the 10X Chromium at the University of Michigan Sequencing Core with a target of 10,000 cells per sample.

#### RNA extraction, cDNA, qRT-PCR

Each analysis includes three biological replicates from three separate differentiation attempts as well as three technical replicates. mRNA was isolated using the MagMAX-96 Total RNA Isolation Kit (Thermo Fisher, Cat#AM1830) or the PicoPure RNA Isolation Kit (Thermo Fisher, Cat#KIT0204). RNA quality and yield was measured on a Nanodrop 2000 spectrophotometer just prior to cDNA synthesis. cDNA synthesis was performed using 100ng RNA per sample with the SuperScript VILO cDNA Kit (Thermo Fisher, Cat#11754250). qRT-PCR was performed on a Step One Plus Real-Time PCR System (Thermo Fisher, Cat#42765592R) using QuantiTect SYBR Green PCR Kit (Qiagen, Cat#204145). Gene expression as a measure of arbitrary units was calculated relative to GAPDH using the following equation:

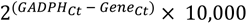

### Bioinformatics/scRNAseq

#### Overview

To visualize distinct cell populations within the single-cell RNA sequencing dataset, we employed the recommended workflow outlined by the Seurat 4.0 R package (Hao *et al*., 2021). This pipeline includes the following steps: filtering cells for quality control, applying the SCTransform technique (Hafemeister and Satija, 2019) in place of traditional log normalization, variable gene selection, and scaling, identifying anchors and integrating if multiple single-cell RNA samples are involved (Stuart *et al*., 2019), reducing dimensionality with principal component analysis (PCA) and uniform manifold approximation and projection (UMAP) (McInnes, Healy and Melville, 2018; Becht *et al*., 2019), clustering by either the Louvain algorithm (Blondel *et al*., 2008) or the Leiden algorithm (Traag, Waltman and van Eck, 2019), and log normalization on RNA assay for final visualization as well as differential gene expression analysis.

#### Sequencing Data and Processing FASTQ Reads into Gene Expression Matrices

All single-cell RNA sequencing was performed at the University of Michigan Advanced Genomics Core with an Illumina Novaseq 6000. The 10X Genomics Cell Ranger pipeline was used to process raw Illumina base calls (BCLs) into gene expression matrices. BCL files were demultiplexed to trim adaptor sequences and unique molecular identifiers (UMIs) from reads. Each sample was then aligned to the human reference genome (hg19) to create a filtered feature bar code matrix that contains only the detectable genes for each sample.

#### Quality Control

To ensure quality of the data, all samples were filtered to remove cells expressing too few or too many genes (Fig. 1E-F/Fig. S1G - <500, >8000; Fig. 2G/Fig. S2C-E/Fig. S3A - <500, >7000; Fig. 3E-H/Fig. S4B - <200, >9500), with too low or too high UMI counts (Fig. 1E-F/Fig. S1G – <500, >50000; Fig. 2G/Fig. S2C-E/Fig. S3A – <500, >50000; Fig. 3E-H/Fig. S4B - <200, >50000), or a fraction of mitochondrial genes greater than 0.1. Following the above steps, a total of (Fig. 1E-F/Fig. S1G – 1067 cells, 36601 genes; Fig. 2G/Fig. S2C-E/Fig. S3A – 9133 cells, 36601 genes; Fig. 3E-H/Fig. S4B – 4334 cells, 36602 genes) were kept for downstream analysis and visualization.

#### SCTransform and Integration

Seurat’s SCTransform method allows efficient pre-processing, normalization, and variance stabilization of molecular count data from scRNA-seq samples. Running this algorithm will reveal a model of technical noise in the scRNA-seq data through “regularized negative binomial regression”, whose residuals are returned as the SCTransform-normalized values that can be used for further downstream analysis such as dimension reduction. During the SCTransform process, we also chose to regress out a confounding source of variation – mitochondrial mapping percentage. When dealing with one sample (Fig. 1E-F/Fig. S1G), there’s no batch effect. But when multiple samples are present (Fig. 2G/Fig. S2C-E/Fig. S3A/Fig. 3E-H/Fig. S4B), we have noticed certain amount of batch effects when clustering data due to technical artifacts such as timing of data acquisition or differences in dissociation protocol. To mitigate these effects, we chose to follow Seurat’s integration workflow due to its optimal efficiency in harmonizing large datasets. After each sample is SCTransform-prepared, a set of anchors, which are mutual nearest neighbors, are identified and filtered. Finally, iterative pairwise integration is performed on the anchor set. After completion of such batch correction, cell clustering should no longer be driven by technical artifacts.

#### Dimension Reduction and Clustering

Principal component analysis (PCA) was conducted on the corrected expression matrix followed. Using the top principal components, a neighborhood graph was calculated for the 20 nearest neighbors (Fig. 1E-F/Fig. S1G – 20 principal components; Fig. 2G/Fig. S2C-E/Fig. S3A – 30 principal components; Fig. 3E-H/Fig. S4B – 30 principal components). The UMAP algorithm was then applied for visualization on 2 dimensions. Using the Leiden algorithm, clusters were identified with a resolution of (Fig. 1E-F/Fig. S1G – 0.2; Fig. 3E-H/Fig. S4B – 0.2). Using the Louvain algorithm, clusters were identified with a resolution of (Fig. 2G/Fig. S2C-E/Fig. S3A – 0.08)

#### Cluster Annotation

Using canonically expressed gene markers, each cluster’s general cell identity was annotated. Markers utilized include epithelium (*EPCAM, KRT18, KRT8, CLDN6*), mesenchyme (*POSTN, DCN, COL1A2, COL3A1*), neuronal (*S100B, STMN2, ELAV4, ASCL1*), endothelial (*ESAM, CDH5, CLDN5, KDR*), proliferative (*MKI67, TOP2A, CDK1*), primordial germ cell (*POU5F1, NANOG, TBXT, NANOS3, TFAP2C*), and foregut mesoderm (*ISL1, HAND1, BMP4, FOXF1, LEF1*).

#### Cell Scoring

Application of cell scoring strategy is as previously described (Holloway *et al*., 2020; Hein *et al*., 2021). Briefly, cells were scored based on expression of a set of 93 – 153 marker genes per cell type. Gene lists were compiled by analyzing previously-published data from human fetal lung (Miller *et al*., 2020; Hein *et al*., 2021). Clusters were first identified by major cell classes (epithelium, mesenchyme, neuronal, endothelial, immune), then the epithelium was sub-clustered to identify bud tip progenitor, basal and neuroendocrine cell clusters by visualizing canonical marker gene expression for each respective cell type. In the case of bud tip progenitor and basal cells, clusters were again sub-clustered to identify the clusters with enriched bud tip progenitor or basal cell marker expression, respectively. The top 50 genes from each bud tip progenitor, basal or neuroendocrine cell clusters were merged to create the gene sets for cell scoring. After obtaining the scaled expression values for the data set, scores for each cell were calculated with the AddModuleScore function of Seurat. Cell scores were visualized by violin plots.

#### Normalization for Visualization and Differential Gene Expression

As recommended by Seurat developers, we employed the method of log normalization on the standard RNA assay for graphing dot plots, feature plots, and conducting DGEs. Expression matrix read counts per cell were normalized by the total expression, multiplied by a scale factor of 10000, and finally log-transformed. For the differential gene expression testing, we only tested features that are first, detected in a minimum fraction of 0.25 in either of the two cell populations, and second, show at least 0.25-fold difference in log-scale between the two cell populations on average.

### Quantification, Statistical Analysis

Graphs and statistical analysis for RT-qPCR and FACS quantification were performed in GraphPad Prism Software. See Figure legends for the number of replicates used, statistical test performed, and the p-values used to determine the significance for each analysis.

### Fluorescent Activated Cell Sorting (FACS) and Flow Cytometry

#### hPSC Flow Cytometry

1mL accutase (Sigma, Cat#A6964) was added to each well of hPSC cultures in a 6-well plate and was incubated at 37°C for 5 – 10 minutes, until cells detached. An equal volume of mTeSR Plus media (StemCell Technologies, Cat#100-0276) with 10µM Y-27632 (APExBIO, Cat#B1293) was added to cells and cells were dissociated mechanically by pipetting with a P1000 pipette 2x, were centrifuged at 300g for 5 minutes at 4°C, excess media was removed, and cells were resuspended in FACS Buffer (2% BSA, 10µM Y-27632, 100U/mL penicillin-streptomycin). Cells were passed through a 70µm cell strainer, pre-coated with FACS buffer, and centrifuged at 300g for 5 minutes at 4°C. Cells were resuspended in 1mL FACS Buffer and transferred to 5mL FACS tubes (Corning, Cat#352063). 0.2µg/mL DAPI was added to respective tubes. Flow cytometry was performed using a Bio Rad Ze5#3 and accompanying software.

#### 3D Culture Sorting

3D cultures (LPOs, iBTOs) were removed from Matrigel using a P1000 pipette tip and vigorously pipetted in a 15mL conical tube to remove as much Matrigel as possible. Tissue was centrifuged at 300g for 3 minutes at 4°C, then excess media and Matrigel was removed. Tissue was digested to single cells using 2 – 4mL TrypLE (Invitrogen, Cat#12605010), depending on pellet size, and incubated at 37°C for 30 minutes, adding mechanical digestion with a pipette every 10 minutes. After 15 minutes, DNase I (Qiagen, Cat#79254) was added to the digestion at 7.5µL/mL TrypLE. After 30 minutes, trypsinization was quenched with DMEM/F-12 (Corning, Cat#10-092-CV) + 10µM Y-27632 (APExBIO, Cat#B1293). Cells were passed through a 70µm cell strainer, pre-coated with DMEM/F-12 +10µM Y-27632, and centrifuged at 500g for 5 minutes at 4°C. Cells were resuspended in 4mL FACS Buffer (2% BSA, 10µM Y-27632, 100U/mL penicillin-streptomycin) and transferred into 5 mL FACS tubes (Corning, Cat#352063). Cells were centrifuged again at 300g for 3 minutes at 4°C, then resuspended in 1mL FACS buffer and counted. 10^5^ cells were placed into new FACS tubes for all controls (no antibody, DAPI only, individual antibodies/fluorophores) and all remaining cells were centrifuged and resuspended in FACS buffer for a concentration of 10^6^ cells/100µL. Primary antibodies were incubated for 30 minutes on ice. 3mL FACS buffer was added to each tube after 30 minutes and tubes were centrifuged at 300g for 3 minutes at 4°C. Cells were washed again with 3mL FACS buffer and centrifuged at 300g for 3 minutes at 4°C. Secondary antibodies were incubated for 30 minutes on ice. 3mL FACS buffer was added to each tube after 30 minutes and tubes were centrifuged at 300g for 3 minutes at 4°C. Cells were washed again with 3mL FACS buffer and centrifuged at 300g for 3 minutes at 4°C. Cells were resuspended in FACS buffer and 0.2µg/mL DAPI was added to respective tubes. FACS was performed using a Sony MA900 cell sorter and accompanying software. Cells were collected in 1mL 3F media +10µM Y-27632. Secondary antibodies were purchased from Jackson Immuno and used at a dilution of 1:500.

## ACKNOWLEDGEMENTS

We would like to thank the University of Michigan Advanced Genomics core for the operation of the 10X Chromium single cell capture platform, the University of Michigan Microscopy core for providing access to confocal microscopes and the Flow Cytometry core for providing access to flow cytometers.

## AUTHOR CONTRIBUTIONS

RFCH, ASC and JRS conceived the study. JRS supervised the research. RFCH, ASC and AF designed, performed, and interpreted experiments optimizing and characterizing NKX2-1^+^ spheroids, LPOs and iBTOs and involving iAirO and iAlvO differentiations. ZX and TF performed computational analysis on scRNA-seq data, and RFCH, ASC and JRS interpreted scRNA-seq results. RFCH, ASC, Y-HT and EMH developed and executed tissue dissociation methods for FACS and scRNA-seq and operated flow cytometers. RFCH, ASC and CJC performed whole mount staining and imaging. SH developed reagents and protocols for hPSC culture and differentiations. JM provided the NKX2-1-EGFP/TP63-mCherry iPSC reporter line and provide insight into the study. RFCH, ASC and JRS assembled figures and wrote the manuscript. ZX contributed to writing the methods. All authors read, contributed feedback, and approved the manuscript.

## DATA AVAILABILTY

Sequencing data used in this study is deposited at EMBL-EBI ArrayExpress. Single-cell RNA sequencing of human fetal lung and human fetal bud tip organoids: human fetal lung (ArrayExpress: E-MTAB-8221, ArrayExpress: E-MTAB-10662) (Miller *et al*., 2020; Hein *et al*., 2021), spheroids, LPOs and iBTOs (in progress) (this study). Code used to process data can be found at: https://github.com/jason-spence-lab/Hein_Conchola_2022.

---

**Supplementary Figure 1:**
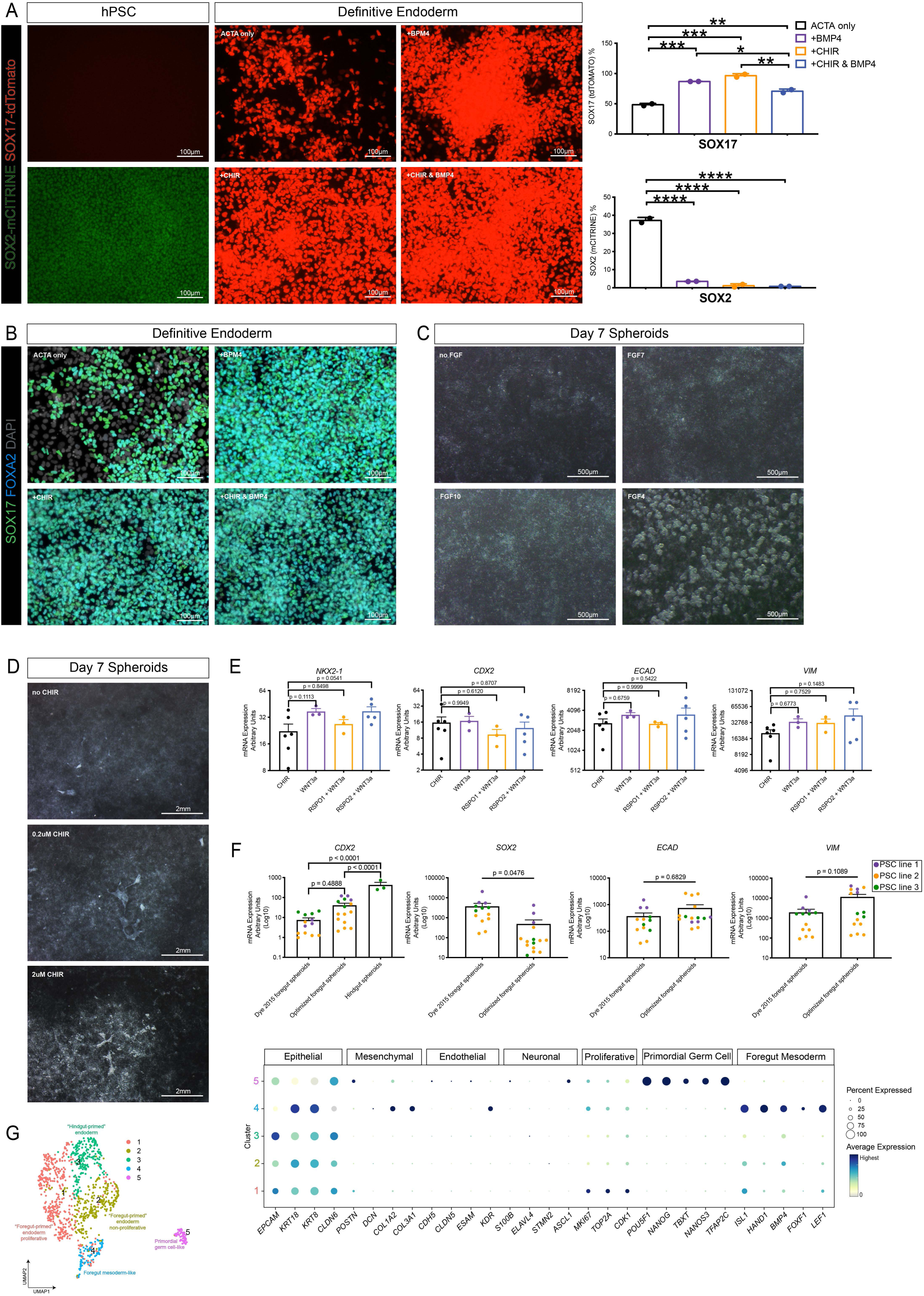
Optimization of Directed Differentiation of hPSCs at Definitive Endoderm, Foregut Spheroid and NKX2-1^+^ Spheroid Stages. (A) SOX2-mCitrine (pluripotent stem cell marker) and SOX17-tdTomato (definitive endoderm marker) reporter images and flow cytometry quantifications of SOX2-mCITRINE and SOX17-tdTomato on hPSCs (images only) and hPSC-derived day 4 definitive endoderm (DE) (images and flow cytometry) where 50ng/mL BMP4 and/or 2uM CHIR were added on day 1 of a 3-day 100ng/mL Activin A (ACTA) treatment to induce DE (see Fig. 1A) in a hPSC line expressing SOX2-mCitrine/SOX17-tdTomato hPSC line (Martyn *et al*., 2018). Two technical replicates from the same experiment were performed for flow cytometry quantifications. Statistical tests were performed by ordinary one-way ANOVA. (B) Immunofluorescence staining for the DE markers SOX17 and FOXA2 on day 4 DE where 50ng/mL BMP4 and/or 2uM CHIR were added on day 1 of a 3-day 100ng/mL Activin A (ACTA) treatment to induce DE (see Fig. 1A). This experiment was performed on two independent hPSC lines different than the reporter line in Fig. S1A. Representative images are shown from H9 ESCs. (C) The requirement of FGF for the formation of spheroids on day 7 of the NKX2-1-optimized spheroid directed differentiation protocol (see Fig. 1A) was tested by adding no FGF, 10ng/mL FGF7, 500ng/mL FGF10, or 500ng/mL FGF4 on days 4 – 6. (D) The requirement of WNT for the formation of spheroids on day 7 of the NKX2-1-optimized spheroid directed differentiation protocol was tested by adding no CHIR99021, 0.2uM CHIR99021, or 2uM CHIR99021 on days 4 – 6. (E) RT-qPCR data comparing expression of the lung epithelial marker *NKX2-1*, the hindgut epithelial marker *CDX2*, the pan-epithelial marker *ECAD*, and the pan-mesenchymal marker *VIM* when CHIR99021, WNT3a, RSPO1 & WNT3a, or RSPO2 & WNT3a was used to activate WNT signaling on days 7 – 9 of the NKX2-1-optimized spheroid directed differentiation protocol. Data points for CHIR99021 and WNT3a + RSPO2 represent 2 independent experiments with 2 – 3 technical replicates and data points from other conditions represent one independent experiment with 3 technical replicates. All experiments were performed on H9 ESCs. Statistical tests compared all conditions to CHIR99021 and were performed by ordinary one-way ANOVA followed by Dunnett’s multiple comparison test. (F) RT-qPCR data comparing expression of the hindgut epithelial marker *CDX2*, the posterior foregut marker *SOX2*, the pan-epithelial marker *ECAD*, and pan-mesenchymal marker *VIM* from previously published foregut spheroids (Dye *et al*., 2015) to optimized foregut spheroids, and in the case of *CDX2*, hindgut spheroids. Each color represents an independent experiment with a unique iPSC line (purple: WTC11, orange: iPSC17 WT 7B2, green: iPSC line 72.3). Each data point of the same color represents a technical replicate from the same iPSC line in one or more independent experiments. Error bars represent standard error of the mean. Statistical test for *CDX2* was performed by ordinary one-way ANOVA followed by Turkey’s multiple comparison test and statistical tests for the remaining markers was performed by a paired t test. (G) Cluster plot of scRNA-seq data from day 10 spheroids and dot plot of epithelial, mesenchymal, endothelial, and neuronal lineage genes as well as proliferation markers in each cluster of the UMAP. The dot size represents the percentage of cells expressing the gene in the corresponding cluster, and the dot color indicates log-normalized expression level of the gene.

**Supplementary Figure 2:**
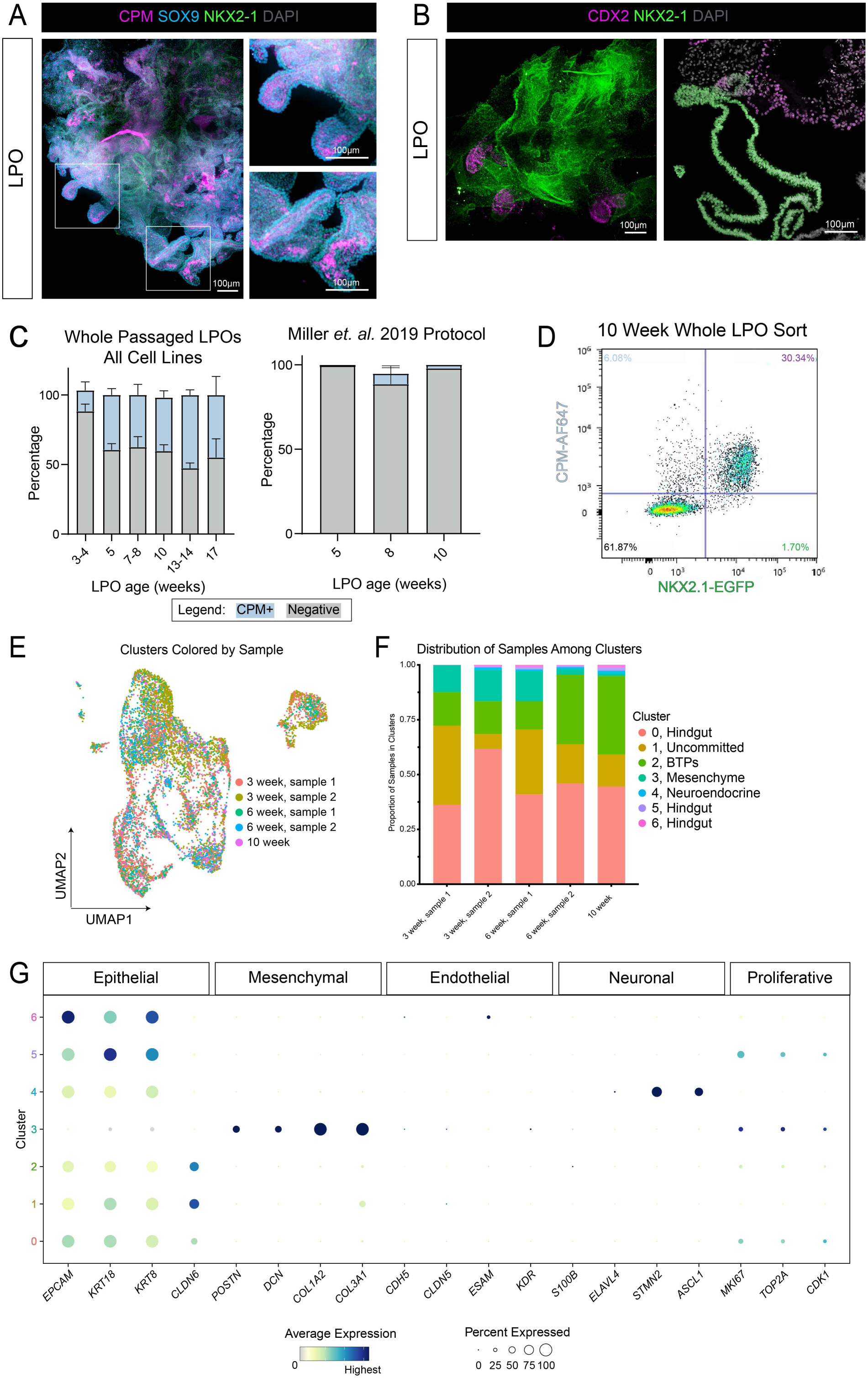
Bud Tip Progenitors and Contaminating Lineages Emerge and Expand in LPOs Over Time. (A) Maximum intensity projection of a whole mount immunofluorescence confocal z-series staining of a 10-week LPO for bud tip progenitor markers CPM and SOX9 and lung epithelial marker NKX2-1. (B) Maximum intensity projection of a whole mount immunofluorescence confocal z-series staining of a 10-week LPO (left) or immunofluorescence staining on paraffin sections of 4-week LPOs (right) for intestinal epithelial marker CDX2 and lung epithelial marker NKX2-1. (C) (Left) FACS quantification of CPM^+^ cells in LPOs in aggregate time course from three separate cell lines, including the NKX2-1-EGFP reporter line, iPSC line 72.3 and WTC11. LPOs were sorted using CPM from 3 – 17 weeks in 3F media. Percentages of live cells expressing CPM (blue) or negative (grey) are reported as mean ± SEM for 2 – 3 replicates per time point. (Right) FACS quantification of CPM^+^ cells in LPOs generated from spheroids using our previous protocol from two separate cell lines, including the NKX2-1-EGFP reporter line and iPSC line 72.3 (WTC11-derived LPOs did not survive to 10 weeks) (Miller *et al*., 2019). (D) Representative flow cytometry plot for iBTP selection based on positive CPM and NKX2-1-EGFP selection on 10-week LPOs. (E) UMAP plot corresponding to the LPO cluster plot in Fig. 2G. Each dot represents a single cell and dots/cells are colored by the sample from which they came from. (F) Stacked bar graph displaying the proportion of cells from each sample in each cluster of the LPO cluster plot in Fig. 2G. (G) Dot plot of epithelial, mesenchymal, endothelial, and neuronal lineage genes as well as proliferation markers in each cluster of the LPO cluster plot in Fig. 2G. The dot size represents the percentage of cells expressing the gene in the corresponding cluster, and the dot color indicates log-normalized expression level of the gene.

**Supplementary Figure 3:**
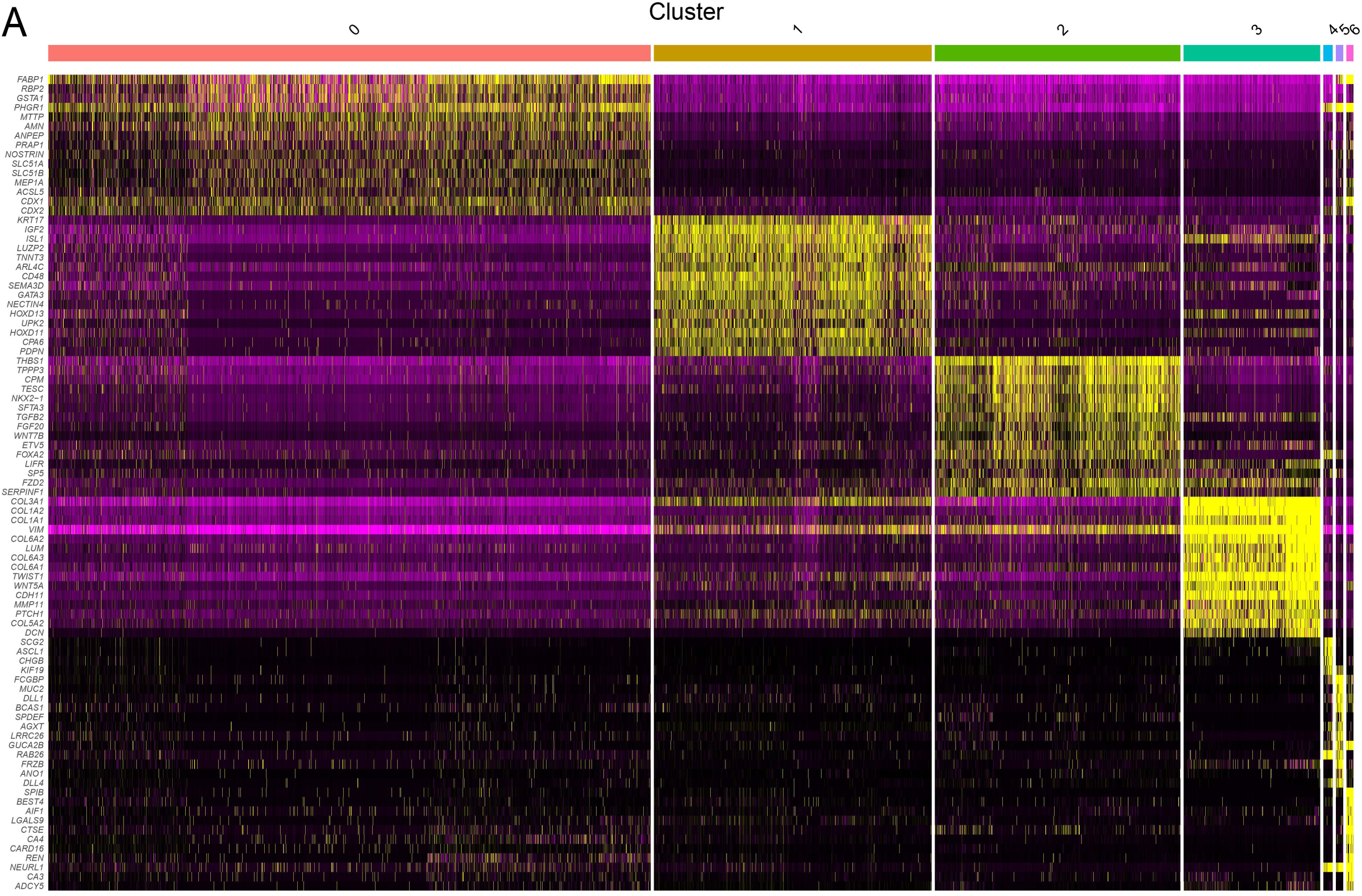
LPOs are Transcriptionally Heterogenous. (A) Heat map including 12 genes per cluster picked from the top-50 highly enriched genes in each cluster corresponding to the UMAP cluster plot in Fig. 2G. Top-50 genes were based on log-fold change of each gene.

**Supplementary Figure 4:**
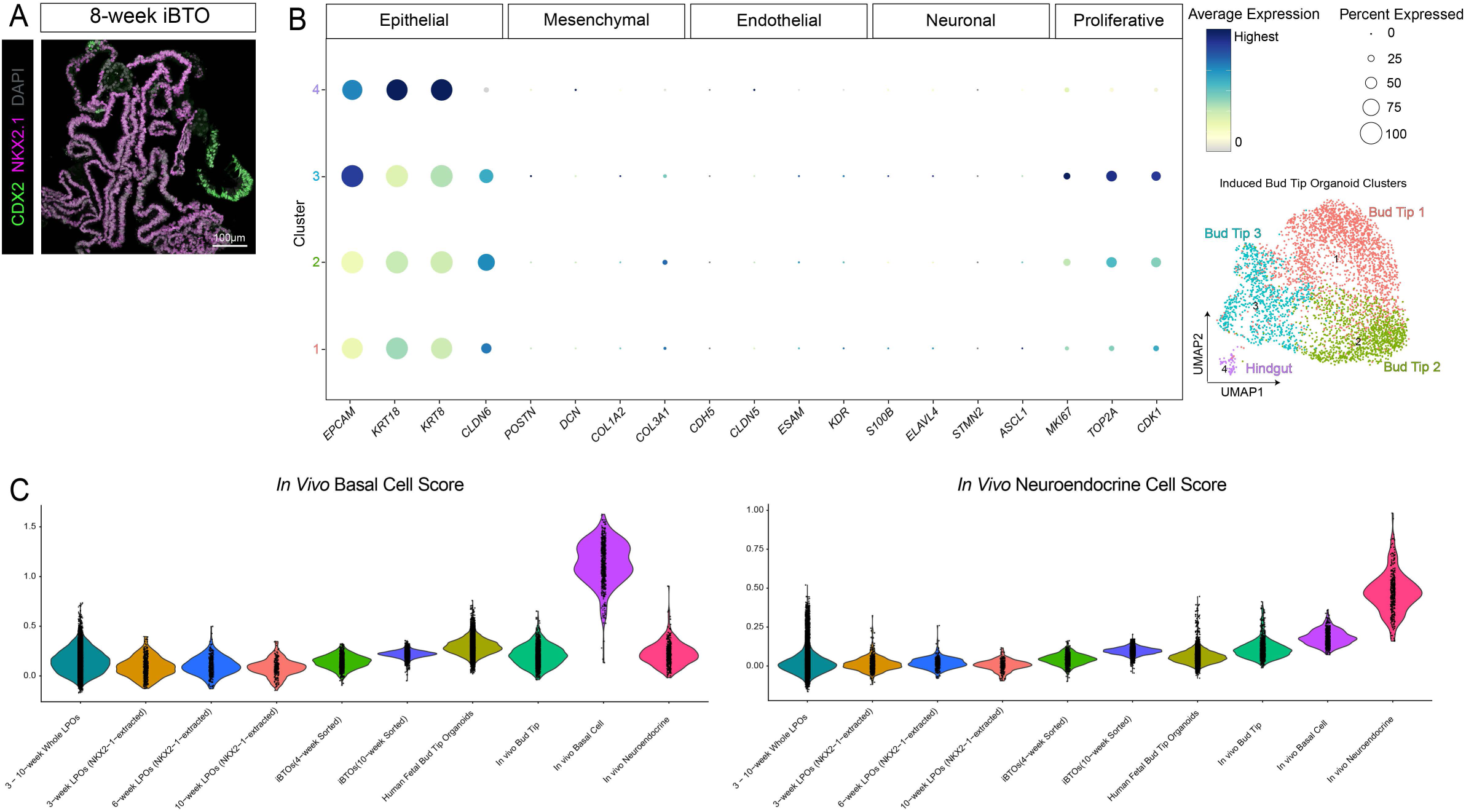
Induced Bud Tip Organoids are Enriched for Bud Tip Progenitor Cells. (A) Immunofluorescence staining on paraffin sections for lung epithelial marker NKX2-1 and intestinal epithelial marker CDX2 on 8-week iBTOs. (B) Dot plot of epithelial, mesenchymal, endothelial, and neuronal lineage genes as well as proliferation markers in each cluster of the iBTO cluster plot in Fig. 3E. The dot size represents the percentage of cells expressing the gene in the corresponding cluster, and the dot color indicates log-normalized expression level of the gene. (C) Violin plot displaying an *in vivo* basal or neuroendocrine cell score, calculated as the average expression of the top 100 (basal) or 150 (neuroendocrine) enriched genes in *in vivo* basal or neuroendocrine cells, for each sample. Samples include whole LPOs (2x3-week LPOs, 2x6-week LPOs, 1x10-week LPOs), NXK2-1-extracted cells from 3-, 6-, and 10-week LPOs (n = 2 for 3- and 6-week LPOs, n = 1 for 10-week LPOs), whole iBTOs (derived from LPOs sorted for NKX2-1^+^/CPM^+^ cells at 4 or 10 weeks, n=1 of each), human fetal-derived bud tip organoids, human fetal-derived bud tip organoids induced to airway (Miller *et al*., 2020) (airway organoids), and primary *in vivo* tissue including computationally-extracted bud tip, basal, and neuroendocrine cells (Miller *et al*., 2020; Hein *et al*., 2021). The cell scoring lists were generated from scRNA-seq data from the *in vivo* basal or neuroendocrine cells shown on the plots.

**Supplementary Figure 5:**
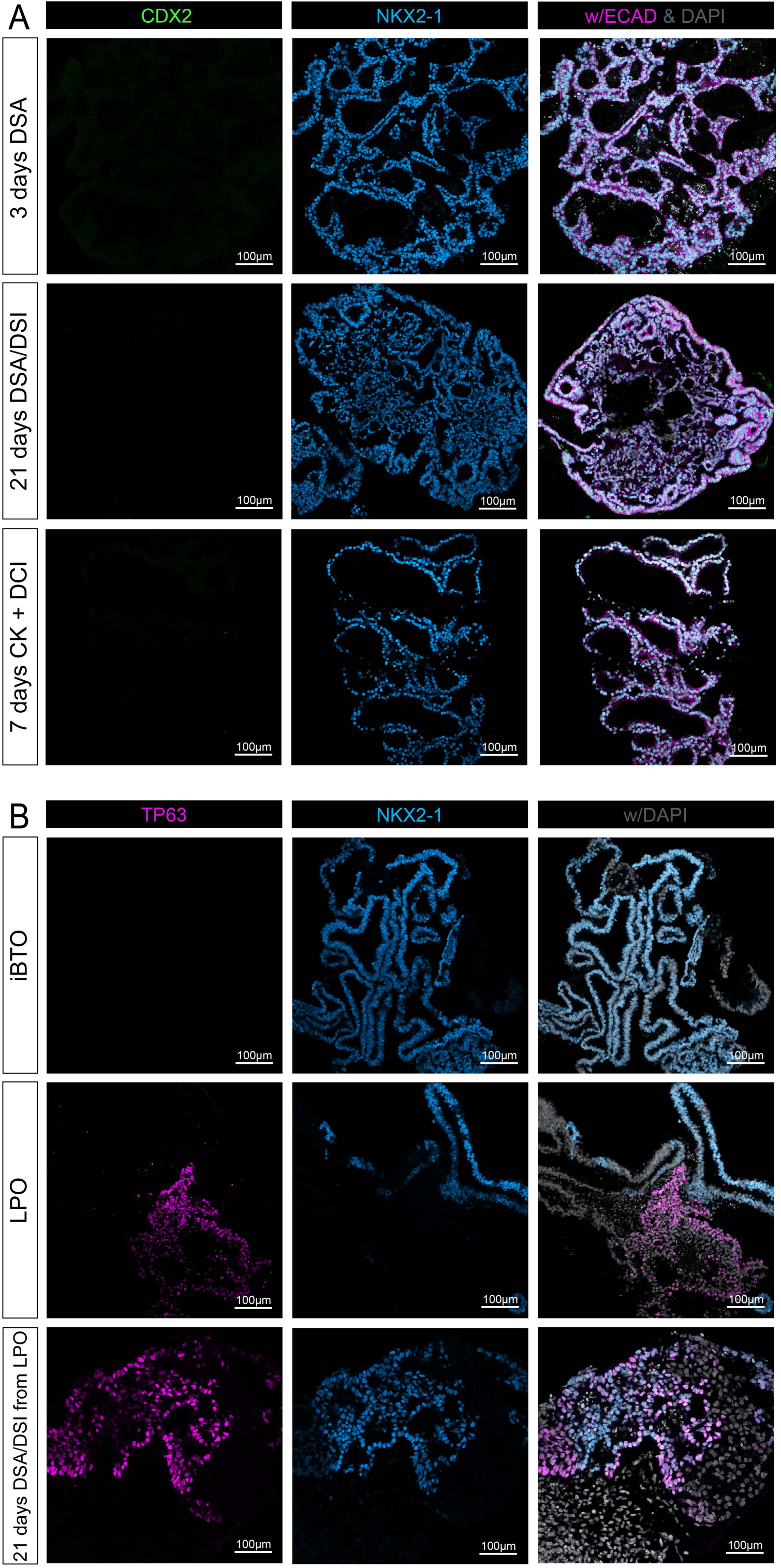
iBTOs Maintain NKX2-1 Expression after Differentiation into Proximal Airway and Distal Alveolar Organoids, and Differentiate More Effectively Compared to LPOs. (A) Immunofluorescence staining on paraffin sections for the intestinal epithelial marker CDX2, lung epithelial marker NKX2-1, and general epithelial cell-type marker ECAD on 12-week iBTOs undergone 3 days of dual-SMAD activation (DSA) or 3 days DSA followed by 18 days of dual-SMAD inactivation (DSI) of the airway induction protocol or 3-week iBTOs undergone the alveolar differentiation protocol (7 days CK + DCI) (see Fig. 4A). (B) Immunofluorescence staining on paraffin sections for the airway progenitor marker TP63 and lung epithelial marker NKX2-1 on 12-week iBTOs, 4-week LPOs, and 3-week LPOs after 21 days of the airway differentiation protocol (DSA/DSI treatment).

